# Methylphenidate alleviates cognitive dysfunction from early Mn exposure: Role of catecholaminergic receptors

**DOI:** 10.1101/2023.06.27.546786

**Authors:** Stephane A. Beaudin, Shanna Howard, Nicholas Santiago, Barbara J Strupp, Donald R Smith

## Abstract

Environmental manganese (Mn) exposure is associated with impaired attention and psychomotor functioning, as well as impulsivity/hyperactivity in children and adolescents. We have shown previously that developmental Mn exposure can cause these same dysfunctions in a rat model. Methylphenidate (MPH) lessens impairments in attention, impulse control, and sensorimotor function in children, but it is unknown whether MPH ameliorates these dysfunctions when induced by developmental Mn exposure. Here, we sought to (1) determine whether oral MPH treatment ameliorates the lasting attention and sensorimotor impairments caused by developmental Mn exposure, and (2) elucidate the mechanism(s) of Mn neurotoxicity and MPH effectiveness. Rats were given 50 mg Mn/kg/d orally over PND 1-21 and assessed as adults in a series of attention, impulse control and sensorimotor tasks during oral MPH treatment (0, 0.5, 1.5, or 3.0 mg/kg/d). Subsequently, selective catecholaminergic receptor antagonists were administered to gain insight into the mechanism(s) of action of Mn and MPH. Developmental Mn exposure caused persistent attention and sensorimotor impairments. MPH treatment at 0.5 mg/kg/d completely ameliorated the Mn attentional dysfunction, whereas the sensorimotor deficits were ameliorated by the 3.0 mg/kg/d MPH dose. Notably, the MPH benefit on attention was only apparent after prolonged treatment, while MPH efficacy for the sensorimotor deficits emerged early in treatment. Selectively antagonizing D1, D2, or α2_A_ receptors had no effect on the Mn-induced attentional dysfunction or MPH efficacy in this domain. However, antagonism of D2R attenuated the Mn sensorimotor deficits, whereas the efficacy of MPH to ameliorate those deficits was diminished by D1R antagonism. These findings demonstrate that MPH is effective in alleviating the lasting attention and sensorimotor dysfunction caused by developmental Mn exposure, and they clarify the mechanisms underlying developmental Mn neurotoxicity and MPH efficacy. Given that the cause of attention and psychomotor deficits in children is often unknown, these findings have implications for the treatment of environmentally-induced attentional and psychomotor dysfunction in children more broadly.

**Highlights:**

- Developmental Mn exposure impaired attention and sensorimotor function
- MPH ameliorated the Mn deficits, but the effective dose varied by functional domain
- The MPH benefit on attention was only apparent after prolonged treatment
- No specific catecholaminergic receptor singly mediates Mn-induced inattention or MPH efficacy in this domain
- Dopamine 1 and 2 receptors moderated Mn sensorimotor deficits and MPH efficacy in this domain
- MPH can improve behavioral dysfunction induced by an environmental toxicant

## 1. INTRODUCTION

Numerous epidemiological studies have demonstrated associations between environmental manganese (Mn) exposure and the prevalence of attention-deficit/hyperactivity disorder (ADHD) and/or attention-impulsivity/hyperactivity symptoms in children and adolescents (Bhang et al., 2013; Bouchard et al., 2007; Broberg et al., 2019; Capelo et al., 2022; Hong et al., 2014; Oulhote et al., 2015). Although suggestive, these studies do not allow causal inferences between Mn exposure and these behavioral impairments due to the possibility that these associations reflect uncontrolled, or poorly controlled, confounding.

Animal model studies are a crucial addition to human investigations of environmental Mn exposure in this regard. First, they are essential for testing whether well-defined Mn exposures and these neurobehavioral symptoms are causally related. Second, they can elucidate the underlying neurobiological bases for the symptoms, which guide treatment options for Mn-exposed children (Smith and Strupp, 2023). Our prior animal model studies have shown that developmental oral Mn exposure at levels corresponding to the exposure risk faced by newborns and young children can produce impaired attention and sensorimotor function, as well as hypofunctioning of the catecholaminergic frontocortico-striatal system – all of which last into adulthood (Beaudin et al., 2017a; Beaudin et al., 2013; Conley et al., 2020; Lasley et al., 2020). These studies have enhanced our understanding of the role of Mn exposure in the development of ADHD symptoms and attentional/sensorimotor deficits in general, and their biological foundations. They also provide a model system for testing the efficacy of potential therapies.

Methylphenidate (MPH) is an effective ADHD medication that can lessen core behavioral symptoms in both children and adults (Coghill et al., 2014; Faraone et al., 2021; Volkow et al., 2005), and also alleviate manual skill impairments in ADHD children with co-morbid developmental coordination disorder (DCD) (Bart et al., 2013). Pre-clinical and clinical studies have implicated a number of mechanisms underlying MPH efficacy, ranging from the acute pharmacologic action of MPH to increase dopaminergic and noradrenergic activity within the fronto-cortico-striatal system (Arnsten, 2009, 2006; Spencer & Berridge, 2019), to longer-term adaptations in catecholaminergic and Wnt- and mTOR-signaling pathways with chronic MPH treatment (Yde Ohki et al., 2020; Zetterström et al., 2022). Overall, however, the specific mechanism(s) responsible for MPH’s efficacy in treating ADHD and related symptoms remain poorly understood.

Our prior studies on the effectiveness of MPH for treating Mn-induced neurobehavioral deficits have demonstrated that 16 days of daily oral treatment at 2.5 mg MPH/kg/d can completely ameliorate Mn-induced impairment in sensorimotor function (Beaudin et al., 2015). However, this MPH regimen did not lessen the attentional dysfunction caused by developmental Mn exposure in adult rats (Beaudin et al., 2017b). These studies, while encouraging, did not identify an MPH dose that can lessen the attentional dysfunction caused by Mn, or provide mechanistic insights into Mn-induced behavioral symptoms and MPH efficacy.

Here, we used our rodent model of developmental Mn exposure to investigate 1) the efficacy of a range of clinically-relevant 30-day oral MPH doses for ameliorating the lasting attention and sensorimotor impairments caused by developmental Mn exposure, and 2) the underlying mechanistic role of dopamine D1, D2, and noradrenergic α2_A_ receptors in developmental Mn neurotoxicity and MPH efficacy. Notably, we used behavioral tests that have been widely employed to assess ADHD-like symptoms and MPH efficacy in rat models of fronto-cortico-striatal dysfunction (Bari et al., 2008; Kloth et al., 2006; Montoya et al., 1991; Robbins, 2002; Whishaw et al., 1997). Our findings demonstrate that MPH is efficacious for treating the lasting attention and sensorimotor deficits caused by developmental Mn exposure, and they clarify the mechanisms underlying developmental Mn neurotoxicity and MPH efficacy.

## 2. MATERIALS AND METHODS

### 2.1. Animals

A total of 128 male Long-Evans rats were used for neurobehavioral evaluation, and an additional 52 male and female littermates were used for blood Mn and hematocrit analyses. Animals were born from 28 nulliparous timed-pregnant rats (Charles River; gestational age 16). Twelve to 24 hours after parturition (designated PND 1, birth = PND 0), each litter was culled to eight pups, with five to six males and the remainder females. Only one male/litter was assigned to a particular treatment condition (See Supplemental Material: *Subjects housing, feeding, and accreditation*).

### 2.2. Study design

The study comprised three successive phases of behavioral evaluation in the same animals: Phase 1 evaluated the lasting neurobehavioral effects of Mn exposure (between-subject); (2) Phase 2 assessed the efficacy of methylphenidate (MPH) treatment (between-subject); and (3) Phase 3 used receptor antagonism (within-subject) to assess involvement of specific catecholaminergic receptors in the Mn deficits and MPH efficacy (see Supplemental Material, *Figure S1 Schematic of study design*). Two Mn exposure levels (0 and 50 mg/kg/d over PND 1-21, n=64 animals/group) were used. The MPH Phase 2 used a 2 x 4 between-subject factorial design, with the animals in each of the two Mn exposure groups assigned to one of the four MPH doses (n = 14-15 animals/group, see *Manganese exposure* and *Methylphenidate treatment* sections below). The final receptor antagonist Phase 3, entailing four receptor antagonist treatments (vehicle, D1R, D2R, α2_A_R), used a within-subject quasi Latin-square design, with each animal receiving all four antagonist treatments (see *Catecholaminergic receptor antagonist treatment* section below).

### 2.3. Manganese exposure

Manganese exposure was carried out as described in our prior studies (Beaudin et al., 2017a; Kern et al., 2010). Neonates were orally exposed to 0 or 50 mg Mn/kg/d from PND 1-21. Manganese was delivered once daily into the mouth of each pup (∼10-25 μL/dose) via a micropipette fitted with a flexible polyethylene pipet tip (Fisher Scientific, Santa Clara, CA, USA). This Mn exposure regimen produces relative increases in Mn intake that approximate the increases reported in infants and young children exposed to Mn via drinking water, diet, or both (See Supplemental Material, *Manganese exposure protocol and rationale*).

### 2.4. Methylphenidate treatment

For study Phase 2, methylphenidate hydrochloride (MPH) (Sigma-Aldrich Inc., St-Louis, MO) was administered orally once daily over a 30-day period from PND 109-143 at doses of 0 (vehicle), 0.5, 1.5, or 3.0 mg/kg/d, as described in our publications (Beaudin et al., 2017b; Beaudin et al., 2015). Control and Mn animals were randomized into the MPH or vehicle treatment groups according to a 2 x 4 between-subject factorial design prior to the start of the MPH phase of the study. MPH was administered daily 30 min before the start of testing using a food wafer delivery method, as previously described (Beaudin et al., 2017b; Beaudin et al., 2015; Ferguson et al., 2009) (See Supplemental Material, *Methylphenidate treatment and rationale*). Because daily testing on the attention task preceded testing on the staircase task, the interval between drug administration and testing was 30 min for the attention tasks, and ∼90 min for the sensorimotor testing. Oral MPH treatment continued into the receptor antagonism Phase 3 of the study, as described below.

### 2.5. Catecholaminergic receptor antagonist treatment with continued MPH treatment

In the antagonist Phase 3 of the study, oral MPH treatment continued while animals also received a subcutaneous injection (1 mL/kg) of one of the catecholaminergic receptor antagonists (SCH23390 for D1R, raclopride for D2R, or BRL 44408 for α2_A_R), or vehicle (saline) 10 minutes before MPH administration and 40 minutes before testing on the selective attention task, and ∼100 min before the staircase task. Antagonist treatments were delivered at doses of 0.005 mg SCH23390/kg s.c., 0.015 mg raclopride/kg s.c., and 1.0 mg BRL 44408/kg s.c. according to a within-subject quasi Latin-square design, in which all rats received all four antagonist treatments in each of three treatment cycles (i.e., three times). To minimize any potential antagonist treatment carry-over effect within a treatment cycle, the antagonist treatments were separated by 2-day washout breaks, with the first washout day consisting of no MPH and antagonist drug treatment and no testing, and the second washout day consisting of MPH treatment only (no antagonist) and testing on the third focused attention task. Each of the three treatment cycles (four antagonist treatment days plus 2 washout days per antagonist) lasted 12 days, and the entire antagonist treatment phase lasted 36 days (PND 144 to 179). Full rationale for the selection of the receptor antagonists, supplier, doses, solution preparation, and timing of treatment are provided in the Supplemental Material (*Catecholaminergic receptor antagonist treatment)*.

### 2.6. Behavioral testing

All animals were tested in the 5-Choice Serial Reaction Time Task (5-CSRTT) and the Montoya staircase test to evaluate learning, attention, impulse control, and sensorimotor functions. Sixteen identical automated 5-CSRTT testing chambers fitted with odor delivery systems (#MED-NP5L-OLF, Med Associates, Inc., St. Albans, VT) were used, as described previously (Beaudin et al., 2017a). Behavioral training began on PND 47, with 1 week of food magazine and nose-poke training, followed by testing on two visual discrimination learning tasks with a fixed cue duration and no pre-cue delay, followed by a series of visual focused and selective attention tasks, as previously described (Beaudin et al., 2017a) (see Supplemental material, *Behavioral testing procedures*).

#### 2.6.1. Focused attention tasks

The first focused attention task used pre-cue delays of 0, 1, 2, or 3 sec and a fixed visual cue duration of 0.7 sec; this task was administered for 12 testing sessions (PND 80-93). The second focused attention task included longer pre-cue delays of 0, 3, 4, or 5 sec and variable visual cue durations of 0.4 or 0.7 sec and was administered for 12 daily testing sessions (PND 95-108). The animals were then tested on a third focused attention task, which included pre-cue delays of 3 or 4 sec and a fixed cue duration 0.7 sec; animals were tested in the third focused attention task for 18 testing sessions/days (PND 109 – 129) while receiving for the first time daily oral MPH treatment.

#### 2.6.2. Selective attention task with olfactory distractors

The subsequent selective attention task was designed to evaluate the rats’ ability to maintain a behavioral or cognitive set in the face of distracting olfactory stimuli (Beaudin et al., 2017a). The selective attention task included the same pre-cue delays and fixed cue duration used in the third focused attention task. In addition, on one third of the trials in each daily testing session, an olfactory distractor was presented from a non-cue response port 1–3 sec before presentation of the visual cue (See Supplemental Material, *Behavioral testing procedures*). Animals were assessed on the selective attention task for 12 testing sessions while they continued to receive daily oral MPH treatment over PND 130-143.

*Dependent measures:* Automatic recorded responses for all attention tasks included %premature responses (responses made after trial onset but before visual cue presentation), %correct response (responses made to the illuminated port following visual cue presentation), %incorrect response (responses made to a non-illuminated port following visual cue presentation), and %omissions (failure to respond within the 10 sec response interval following visual cue presentation). Details about the calculation of response outcomes and other response latency measures are provided in the Supplemental Material (*5-CSRTT dependent measures*).

#### 2.6.3. Staircase apparatus and procedure

Eight Plexiglas staircase devices were used to assess forelimb sensorimotor reaching and grasping skills, as described previously (Beaudin al., 2013). The staircase test procedure included habituation and training over PND 95 - 108, and daily testing over PND 109-129 using a colored-coded food pellet procedure, as previously described (Beaudin et al., 2015; Beaudin et al., 2013).

*Dependent measures*: Forelimb sensorimotor function was measured step-by-step for the number of pellets taken, the number of pellets eaten, the %success, and the number of pellets misplaced. Specifics about the calculation of response outcomes are described in Supplemental Material (*Staircase test dependent measures*).

### 2.7. Blood Mn and hematocrit levels

Whole blood was collected into heparinized containers from littermates to the behaviorally tested animals and analyzed for Mn concentrations (PND 24 and 66 animals) and hematocrit (PND 24), as previously described (Beaudin et al., 2017a; Kern et al., 2010) (see Supplemental Material, *Methods for determining blood Mn concentrations and hematocrit*).

### 2.8. Statistical methods

The behavioral data were modeled by way of structured covariance mixed models (SAS version 9.4 for Windows) and according to the factorial design of each study phase (see *Study Design* section above). Fixed treatment effects included Mn exposure (2 levels corresponding to the two exposure treatment groups), MPH treatment (four levels corresponding to the four MPH-dose groups), and/or antagonist treatment (four levels corresponding to the four receptor antagonist treatments), depending on the study phase. Furthermore, the model included the within-subject factors pre-cue delay, cue duration, day, testing session block, distraction condition, and/or staircase step level, depending on the outcome analyzed. In all models, the random effect was rat to account for correlations within observations from the same animal. The significance level was set at *p* ≤ 0.05, and *p*-values between 0.05 and 0.10 were considered trends and are reported if the pattern of findings aids in clarifying the nature of the Mn, MPH, or antagonist treatment effects. Significant main effects or interaction effects were followed by single-degree of freedom contrasts to clarify the nature of the effects, using the Student’s *t*-test for pairwise comparisons of least squared means. Blood Mn and hematocrit data were analyzed using mixed model analysis of variance and Tukey’s *post hoc* test for pairwise comparisons.

## 3. RESULTS

We focus on the outcome measures which most clearly revealed the effects of Mn exposure and MPH treatment; namely, percent accurate and premature responses for the attention task series, and the number of pellets taken and misplaced for the staircase test of sensorimotor function.

### 3.1. Developmental Mn exposure causes long-term impairments in attention and learning

#### 3.1.1. Manganese exposure impairs focused attention

The ability to maintain attentional focus to brief visual cues presented randomly in space (i.e., response port) and time is impaired by developmental Mn exposure. The analysis of %accurate responses in the first focused attention task reveals a significant interaction between Mn exposure and testing session block (3 sessions per block) (F(3, 434)=6.75, p=0.0002), reflecting that performance of the Mn group improves less across testing blocks than that of controls (Figure 1A); they were also impaired relative to controls in all four blocks (all p’s<0.05). In the second, more challenging focused attention task, an interaction emerges between Mn exposure and pre-cue delay (F(3, 520)=3.67, p=0.012), reflecting that attentional accuracy in the Mn rats, in contrast to controls, did not significantly improve at the longer delays relative to the 0 sec delay. It is notable, however, that the Mn animals were impaired at all pre-cue delays relative to controls (all p’s<0.05) (Figure 1B).

**Figure 1A-D.**
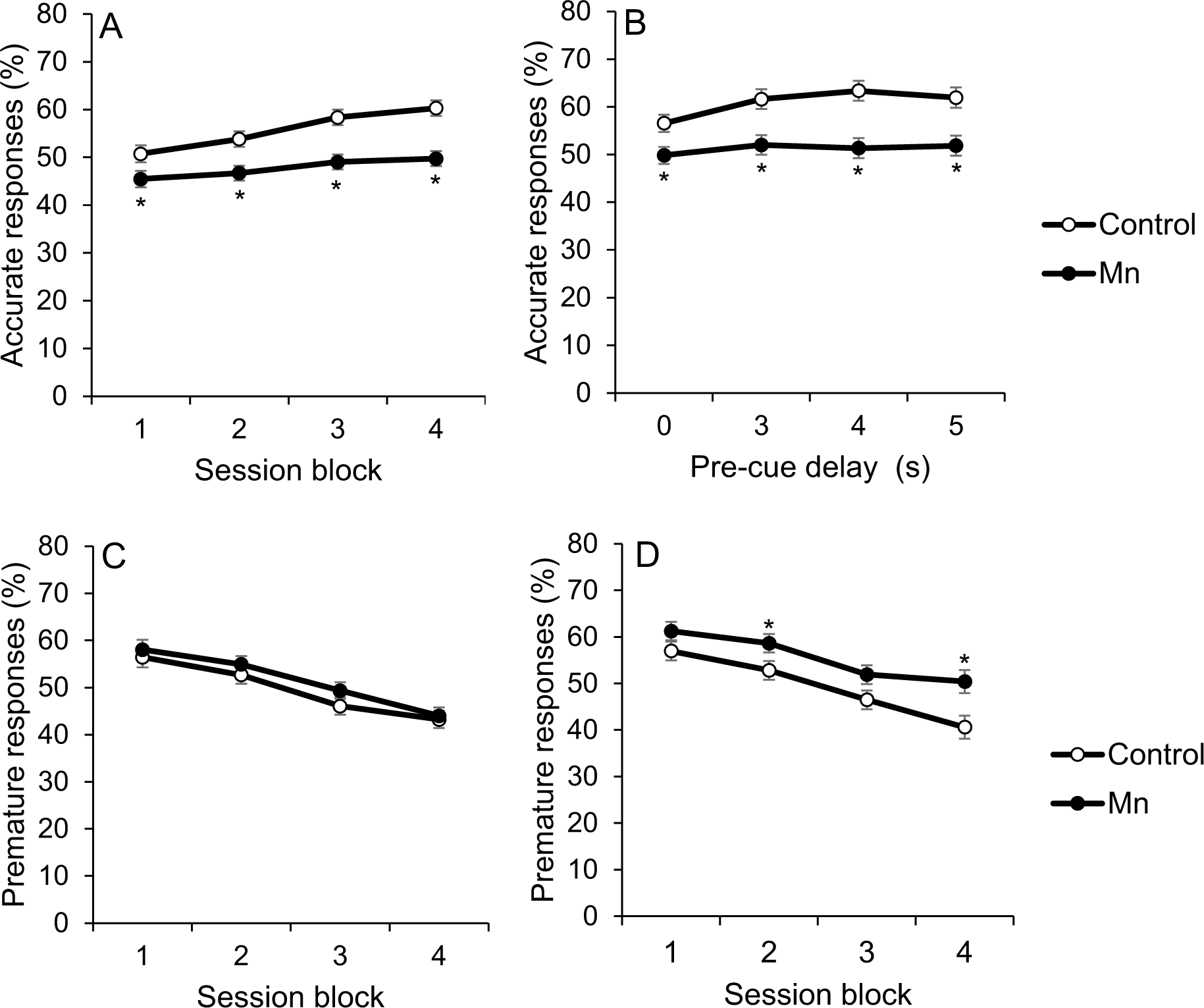
Developmental postnatal Mn exposure produces deficits in attention and learning in the focused attention tasks with variable pre-cue delays. Accurate responses (%) for the control and the Mn groups, as a function of (A) test session block in the first focused attention task, and (B) increasing pre-cue delay in seconds (s) in the second focused attention task. Test session blocks are three successive daily test sessions per block. Premature responses (%) for the control and the Mn groups, as a function of test session block in (C) the first, and (D) second focused attention tasks. Data are lsmeans ± SEM of the control and Mn groups (*n* = 61–63/group). *p≤ .05 versus controls.

#### 3.1.2. In the second focused attention task, Mn exposure impairs learning to withhold responses until the cue was presented

While the %premature responses (i.e., a nose-poke into a port prior to presentation of the visual cue) did not differ between the Mn and control groups in the first focused attention task (Figure 1C), impairment in inhibitory control emerges in the second focused attention task with the imposition of longer pre-cue delays (Figure 1D). Specifically, the analysis of %premature responses reveals a borderline interaction between Mn exposure and testing session block (F(3, 376)=2.22, p=0.085), reflecting that the Mn group learned to withhold premature responses more slowly than controls (Figure 1D). As a result, the Mn group committed significantly more %premature responses than controls during session blocks 2 and 4 (p’s=0.048 and 0.006, respectively) (Figure 1D).

### 3.2. Prolonged MPH therapy at the lowest MPH dose ameliorates Mn-induced attention impairments and improves inhibitory control

Our prior studies using a selective attention task revealed that rats exposed to Mn during early development are more disrupted than controls by unpredictable olfactory distractors, manifested as reduced response accuracy and increased premature responses on distraction trials (Beaudin et al., 2017a; Beaudin et al., 2017b). To determine whether MPH treatment alleviates these areas of dysfunction, animals were dosed with oral MPH and tested on this selective attention task, and a preceding ‘baseline’ task that was identical in terms of cue onset/duration but did not include olfactory distractors (see Methods); this baseline task is referred to as the third focused attention task.

#### 3.2.1. Mn exposure impairs attention in the third focused attention task, and prolonged low dose MPH therapy ameliorated this Mn-induced attentional dysfunction

The major findings in the third focused attention task are: (1) Mn exposure impairs attentional accuracy, (2) oral 0.5 mg MPH/kg/d fully ameliorates the Mn-induced attentional dysfunction, but only after prolonged MPH treatment, and (3) the two higher MPH doses had little to no benefit (relative to vehicle treatment) in ameliorating attentional dysfunction induced by Mn. These findings are based on the significant Mn x MPH x session block interaction for %accuracy (F(15, 747)=4.82, p<0.0001). Specifically, the Mn+0 MPH group was significantly less accurate than unexposed controls over all testing session blocks (p’s<0.001) (Figure 2A). Notably, for the Mn group treated with 0.5 mg MPH/kg/d, there is little or no MPH efficacy during the initial ∼9d of testing and MPH treatment (session blocks 1 – 3), but complete efficacy by session block six (i.e., session days 16-18), in which the Mn-exposed animals treated with this MPH dose perform like controls given the same MPH dose (p=0.78) (Figure 2A). Moreover, contrasts show that the Mn+0.5 MPH group performed significantly better than the Mn+0 MPH group during session blocks 4–6 (p’s=0.03, 0.01, and 0.04), and was not different than the Control+0 MPH group during session blocks 3-5 (p’s=0.21, 0.13, 0.38) (Figure 2A). In contrast, the higher MPH doses of 1.5 and 3.0 mg/kg/d showed little or no efficacy, i.e., attentional accuracy of the Mn+1.5 MPH group and the Mn+3.0 MPH group was not different from the Mn+0 MPH group and was significantly worse than the Control+0 MPH group across many session blocks (all p’s<0.05) (Figure 2A). Finally, we also evaluated the MPH dose-response function for attentional accuracy and found that it is substantially different in the Mn animals (inverted U-shaped) compared to the control animals (U-shaped), further supporting fundamental neurobiological differences between Mn and control animals (see Supplemental, *Figure S2*).

**Figure 2A-D.**
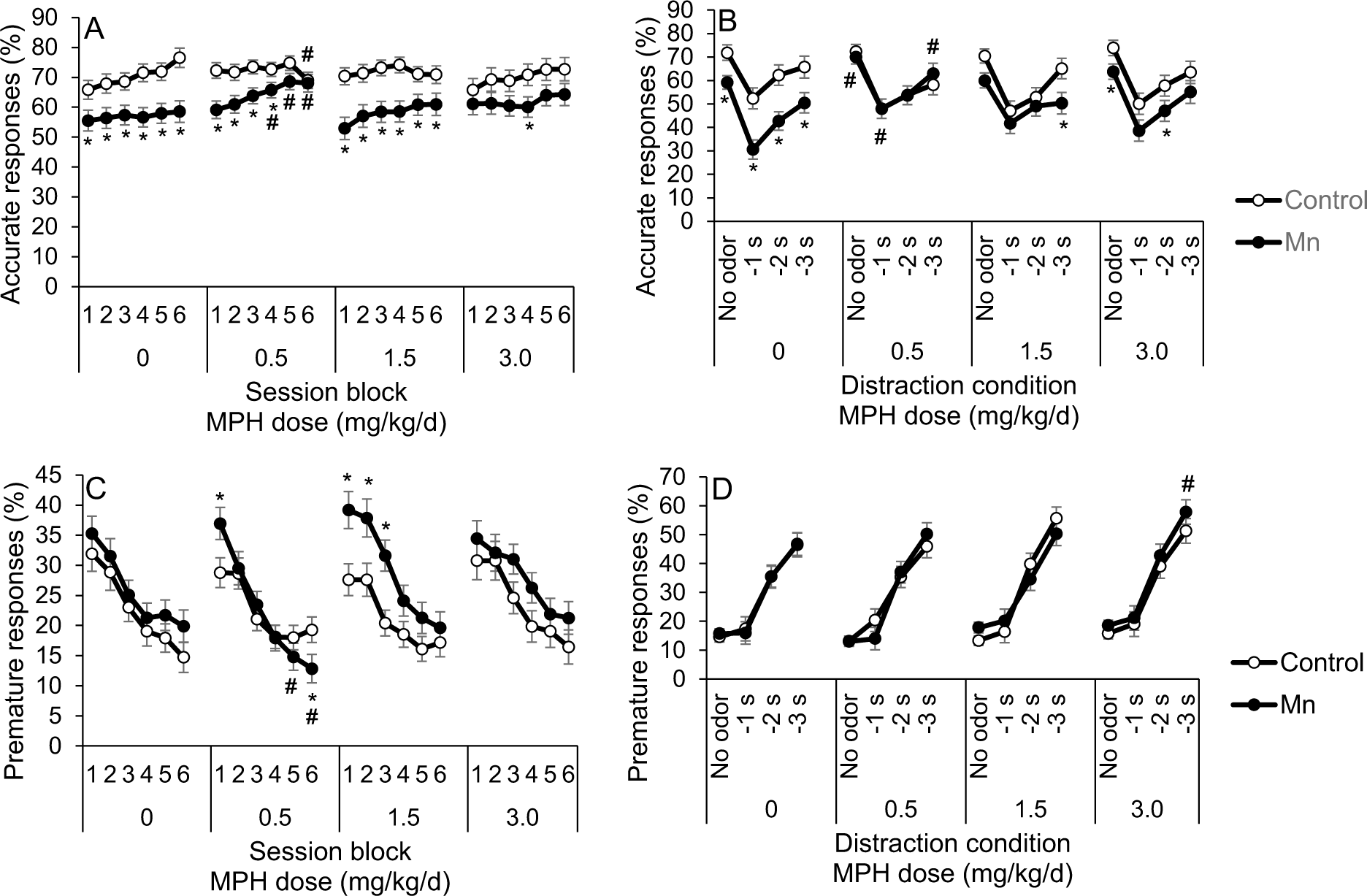
**Panels A, B**. Prolonged 0.5 mg/kg/d methylphenidate (MPH) treatment alleviates Mn attention deficits in the third focused attention task and the selective attention tasks with distractors. (A) Accurate responses (%) for the control and Mn groups, as a function of test session block and MPH dose in the third focused attention task. (B) Accurate responses (%) for the control and the Mn groups, as a function of distractor and MPH dose in the selective attention task. **Panels C, D**. Chronic low dose methylphenidate treatment decreases impulsivity in the Mn-exposed rats in the third focused attention task but heightens impulsivity at the highest MPH dose in the selective attention task. (C) Premature responses (%) for the control and the Mn groups, as a function of test session block and MPH dose in the third focused attention task. (D) Premature responses (%) for the control and the Mn groups, as a function of distractor and MPH dose in the selective attention task. For panels A and C, each test session block contains three daily test sessions. Data are lsmeans ± SEM of the control and Mn groups (*n* = 14–15/group). \**p*≤0.05 versus controls and ^#^*p*≤0.05 versus 0 mg MPH/kg/d dose.

#### 3.2.2. The selective attention task with olfactory distractors reveals continued attentional dysfunction in the Mn animals versus controls, and importantly, that the 0.5 mg/kg/d MPH dose completely ameliorates this dysfunction (*Figure 2C*)

The inferences that the Mn animals show continued attentional dysfunction in the selective attention task with olfactory distractors, and that the 0.5 mg/kg/d MPH dose completely ameliorates this dysfunction are based on the results of the analysis of percent accuracy, which revealed a significant Mn x MPH x distraction condition interaction (F(9, 558)=2.18, p=0.022). This interaction reflects, in part, that although the Mn+0 MPH group exhibit significantly lower percent accurate responses than the Control+0 MPH group in all distraction conditions (p’s<0.05), the Mn impairment is most apparent when the olfactory distractor is presented 1 sec before the visual cue (i.e., the −1s condition) – the condition under which the olfactory stimulus most disrupts performance in both groups. This pattern indicates an impaired ability to maintain attentional focus, which is exacerbated by distractors (i.e., impairments in both focused and selective attention) (Figure 2C).

The significant 3-way interaction of Mn exposure, MPH dose, and distraction condition also reflects that MPH treatment at the 0.5 mg/kg/d dose (but not higher doses) fully ameliorates the Mn attentional dysfunction in this task. Specifically, the Mn+0.5 MPH group performed identically to the Control+0.5 MPH and Control+0 MPH groups under all distraction conditions, and significantly better than the Mn+0 MPH group (p’s=0.01, 0.003, 0.055, and 0.041 for the no odor, −1, −2, and −3 s distraction conditions, respectively) (Figure 3A). Also, the Mn+0.5 MPH group performed indistinguishably from the Control+0 MPH group across all distraction conditions (p’s=0.69, 0.47, 0.15, and 0.66) (Figure 3A). In contrast, the higher MPH doses produced intermediate or no improvement in attentional accuracy in the Mn animals. For example, attentional accuracy of the Mn+1.5 MPH and Mn+ 3.0 MPH groups was not significantly different than the Mn+0 MPH group for any distraction condition; although for some distraction conditions, the Mn animals treated with either the 1.5 or 3.0 MPH dose were also not different from controls (Figure 3A). This latter finding suggests an intermediate efficacy of these higher MPH doses for the Mn animals.

**Figure 3A, B.**
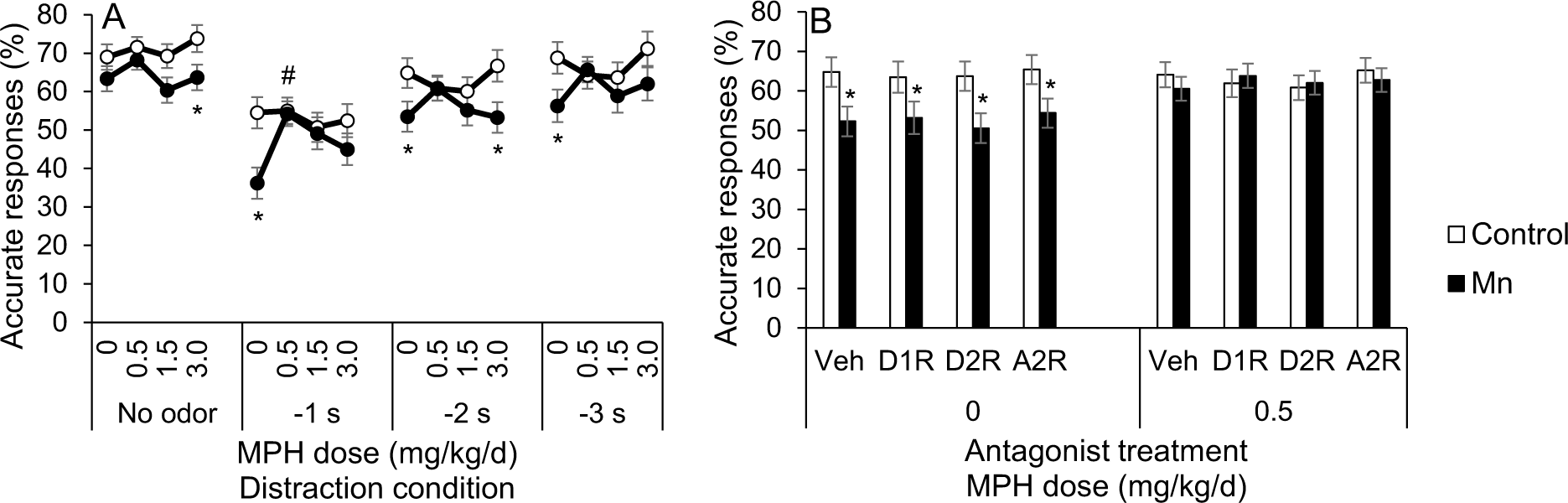
The attention deficit induced by Mn and MPH efficacy to alleviate the Mn dysfunction at the lowest 0.5 mg/kg/d dose are sustained into the antagonist treatment phase in the selective attention task, but are not altered by specific D1R, D2R, or α2_A_R antagonists. (A) Accurate responses (%) for the control and the Mn groups, as a function of MPH dose and distraction condition. (B) Accurate responses (%) for the control and the Mn groups, as a function of antagonist treatment and MPH dose. Data are lsmeans ± SEM of the control and Mn groups (*n* = 14–15/group). *p≤0.05 versus controls, and ^#^p≤0.05 versus the 0 mg/kg/d MPH dose. Data in A are lsmeans ± SEM of the Mn and control groups under the vehicle antagonist treatment.

#### 3.2.3. In the third focused attention task, 0.5 mg/kg/d MPH treatment improves impulse control in the Mn-exposed rats

Analysis of %premature responses in the third focused attention task reveals a significant Mn x MPH x testing session block interaction (F(15, 576)=2.41, p=0.002). This interaction reflects that the lowest dose of MPH is effective in improving impulse control in the Mn group, whereas the higher doses are not, and for this low MPH dose the benefit emerged during session blocks 5 and 6 (i.e., p’s=0.041 and 0.050 versus vehicle treatment, respectively, and in block 6 p=0.033 vs the Control+0.5 MPH group; Figure 2C). Finally, as with attentional accuracy, the Mn animals exhibit an inverted U-shaped MPH dose-response function for %premature responses, whereas controls exhibit a U-shaped MPH dose-response function, further supporting that the Mn animals respond differently to MPH than controls (see Supplemental, *Figure S2*).

#### 3.2.4. In the selective attention task, prolonged 3.0 mg/kg/d MPH treatment increases impulsivity in the Mn rats

As shown in our prior studies (Beaudin et al., 2017a; Beaudin et al., 2017b), the presentation of olfactory distractors prior to the visual cue significantly increases premature responding in both control and Mn groups (F(3, 399)=541.87, p<0.0001), particularly under the more challenging response/impulse control conditions where the interval between the presentation of the distractor and visual cue was longer (2 and 3 sec; p’s<0.0001) (Figure 2F). A three-way interaction between Mn, MPH, and distraction condition was also found (F(9, 506)=2.55, p=0.007). This interaction reflects that, while there was no effect of early postnatal Mn exposure on impulse control, the highest dose of MPH (3.0 mg/kg/d) increased premature responding of the Mn animals (relative to the Mn+0 MPH group) specifically for trials with the distractor presented 3 sec before cue onset (p=0.050) (Figure 2F) This latter finding indicates that the Mn animals responded differently than controls to MPH following prolonged treatment with the 3.0 mg/kg/d dose (Figure 2F).

### 3.3. Specific D1R, D2R, or α2_A_R involvement in Mn-induced attention deficit, impulse control, and MPH efficacy

#### 3.3.1. Mn-induced attentional dysfunction and MPH therapeutic efficacy are maintained throughout the catecholaminergic receptor antagonist study phase

To confirm that the Mn-induced attention deficit and MPH efficacy persisted into the 36-day receptor antagonist phase of the study, we first assessed attentional accuracy in control and Mn animals treated with the receptor antagonist vehicle. Results show that both the Mn attentional dysfunction and the efficacy of the 0.5 mg/kg/d MPH dose persisted into the receptor antagonist treatment phase of the study. Specifically, attentional accuracy in the Mn+0.5 MPH group was significantly better than the Mn+0 MPH group (p=0.0008) and was indistinguishable from the control+0 MPH group (p=0.98) at the −1 s distraction condition (Figure 3A).

#### 3.3.2. Mn-induced attentional dysfunction and the efficacy of MPH to ameliorate this area of dysfunction are not singly mediated by catecholaminergic D1, D2, or α2_A_ receptors

Given the efficacy of the 0.5 mg MPH/kg/d dose to fully ameliorate the Mn attention deficits in the selective attention task, we focused subsequent analyses on this MPH dose to determine whether D1R, D2R, or α2_A_R antagonism altered the Mn attentional dysfunction or MPH efficacy. The results show that none of the specific D1R, D2R, or α2_A_R antagonists altered the Mn-induced attentional dysfunction or the efficacy of the lowest MPH dose to improve attention. The analysis uncovered a significant 3-way interaction of Mn, MPH, and distraction condition (F(3, 212)=3.32, p=0.021), but no significant interactions involving Mn, MPH, and antagonist treatment (F(3, 670)=0.97, p=0.41). Specifically, in the absence of MPH treatment, the Mn-exposed animals (Mn+0 MPH) performed significantly worse than their unexposed counterparts (Control+0 MPH; p=0.021), and this Mn impairment remained under D1R, D2R, and α2_A_R antagonist treatment (p’s=0.070, 0.015, and 0.036, respectively) (see Figure 3B). Furthermore, attentional accuracy of the Mn+0.5 MPH group was indistinguishable from that of the control+0 MPH group across all receptor antagonist treatment conditions (Figure 3B). We conclude from these findings that neither the D1, D2, or α2_A_ receptor is singly driving the Mn-induced attentional dysfunction, or the therapeutic efficacy of MPH treatment for this area of dysfunction.

#### 3.3.3. D1R and D2R antagonists influence impulsivity and correct response latency under certain Mn/MPH treatment conditions

The effect of the 3.0 mg/kg/d MPH dose to increase premature responses in the Mn rats under the −3s odor distractor condition in the selective attention task (Figure 2F, reported above) was not sustained into the antagonist treatment phase of the study. Premature responses in the Mn+3.0 MPH group did not significantly differ from the Mn+0 MPH group for any distraction condition (p’s>0.05), and the Mn+3.0 MPH group exhibited a similar level of premature responses compared to the control+0 MPH group across distraction conditions (p’s>0.05) (Figure 4A).

**Figure 4A-C.**
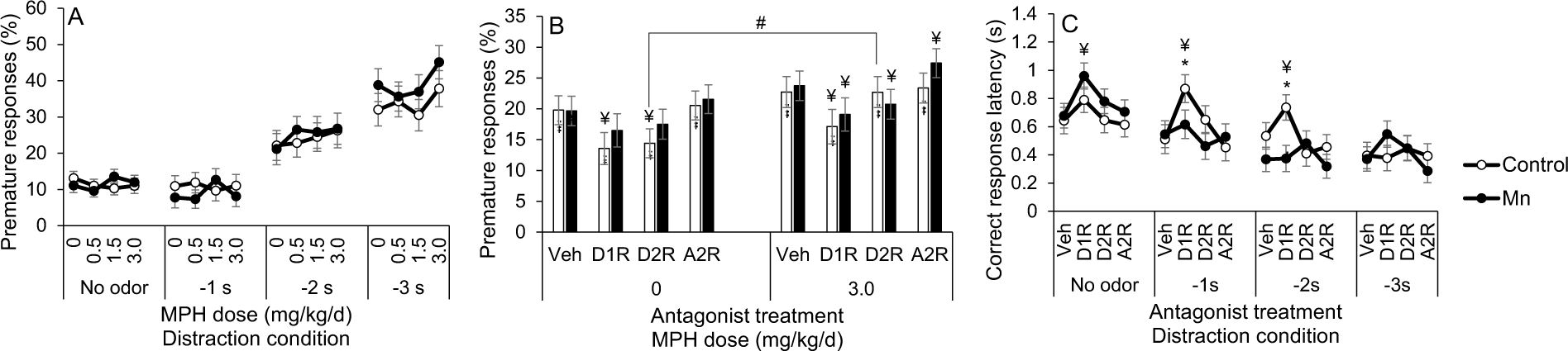
D1R or D2R antagonist treatment decreases premature responses and increases correct response latency, depending on MPH dose and Mn exposure level. (A) Premature responses (%) for the control and the Mn groups, as a function of MPH dose and distraction condition. (B) Premature responses (%) for the control and the Mn groups, as a function of antagonist receptor treatment and MPH dose. (C) Correct response latency for the control and the Mn groups, as a function of antagonist treatment and distraction condition. Data are lsmeans ± SEM of the control and Mn groups (*n* = 14–15/group). *p≤0.05 versus controls, ^#^p≤0.05 versus 0 mg/kg/d MPH dose, and ^¥^p≤0.05 versus vehicle antagonist treatment within MPH condition. Data in A are lsmeans ± SEM of the control and Mn groups under the vehicle antagonist treatment, while data in C are lsmeans ± SEM of the control and Mn groups under the 3.0 mg/kg/d MPH dose.

Nonetheless, due to this prior effect of the 3.0 mg/kg/d MPH dose on impulse control in the Mn group in the selective attention task, we conducted analyses to determine whether any of the antagonists altered impulse control in the control and Mn groups treated with this 3.0 mg/kg/d MPH dose. We also examined correct response latency (i.e., the speed of correct responses), because a reduction in the propensity for premature responses produced by dopamine receptor antagonists has been reported to be accompanied by an increase in response latency due to a delay in the initiation of responding (Harrison et al., 1997).

The results reveal a borderline interaction of Mn, MPH, and antagonist for percent premature responses (F(3, 590)=2.34, p=0.072), reflecting different effects of the catecholaminergic receptor antagonists depending on the Mn exposure and MPH dose. First, the effect of the D2R antagonist on premature responses in control rats varied as a function of MPH dose. For example, for control rats not treated with MPH, the D2R antagonist significantly reduced premature responses compared to the vehicle (p=0.0006), while having no effect on controls chronically treated with 3.0 MPH versus their respective vehicle (p=0.99) (Figure 4B). Moreover, the control+3.0 MPH rats treated with the D2R antagonist had significantly higher premature responses compared to their Control+0 MPH counterparts treated with the D2R antagonist (p=0.02) (Figure 4B). In contrast, the effects of the D2R antagonist in the Mn-exposed rats exhibited a different pattern. For the Mn+0 MPH animals, the D2R antagonist had no impact on premature responding (compared to their Mn+0 MPH+Veh counterpart, p=0.19), but significantly reduced impulsivity in the Mn animals chronically treated with 3.0 MPH (versus their Mn+3.0 MPH+Veh counterpart, p=0.05) (Figure 4B).

The analysis of correct response latency for animals treated chronically with the 3.0 MPH dose revealed a significant Mn x antagonist x distraction condition interaction (F(9, 619)=2.16, p=0.023) (Figure 4C). This interaction reflects that the correct response latency was significantly increased by the D1R antagonist (vs. vehicle) under the no odor condition for the Mn group (p=0.009), but not for controls (p=0.18) (Figure 4C). This contrasts sharply with the D1R antagonist’s effect on increasing latencies to initiate a correct response (versus vehicle) in the control group when a distractor was presented −1 sec or −2 sec before presentation of the visual cue (p’s=0.001 and 0.067, respectively), effects that were not observed in the Mn group (p’s=0.52 and 0.96) (Figure 4C). As a result, when animals were treated with the D1R antagonist in the presence of the olfactory distractors, the Control+3.0 MPH group took longer to initiate a correct response than the Mn+3.0 MPH group (p’s=0.076 and 0.006 for the −1 sec or −2 sec condition, respectively). Based on these findings, we can conclude that the 30 days of oral 3.0 mg MPH/kg/d treatment that preceded the receptor antagonist phase of the study altered the number and/or binding sensitivity of the D1 and D2 receptors in the Mn group.

### 3.4. Mn exposure causes lasting impairments in sensorimotor reaching and grasping skills, and these deficits are ameliorated by treatment with MPH

Developmental Mn exposure caused lasting deficits in sensorimotor skills to reach and grasp food pellets in the Montoya staircase task, and both types of dysfunction are ameliorated by the higher doses of MPH. This is shown by the significant Mn x MPH x step interaction for the number of pellets taken (F(15, 1552)=5.17, p<0.0001) and the number of pellets misplaced (F(15, 1614)=2.95, p=0.0001). The 3-way interaction for pellets taken reflects that the Mn+0 MPH group took significantly fewer pellets than their Control+0 MPH counterparts at steps 5 and 6 (p’s=0.024 and 0.0041, respectively), and that this Mn deficit was ameliorated by the 1.5 and 3.0 MPH doses (Figure 5A). Moreover, the Mn+3.0 MPH group retrieved significantly more pellets than the Mn+0 MPH group at staircase step 6 (p=0.05), and a similar number of pellets than the Control+0 MPH group at the same step (p=0.33) (Figure 5A). In contrast, the lower 0.5 mg MPH/kg/d dose had no efficacy in alleviating this area of dysfunction; i.e., the number of pellets taken by the Mn+0.5 MPH group was not different from the Mn+0 MPH group at step 6 (p=0.51) (Figure 5A).

**Figure 5A, B.**
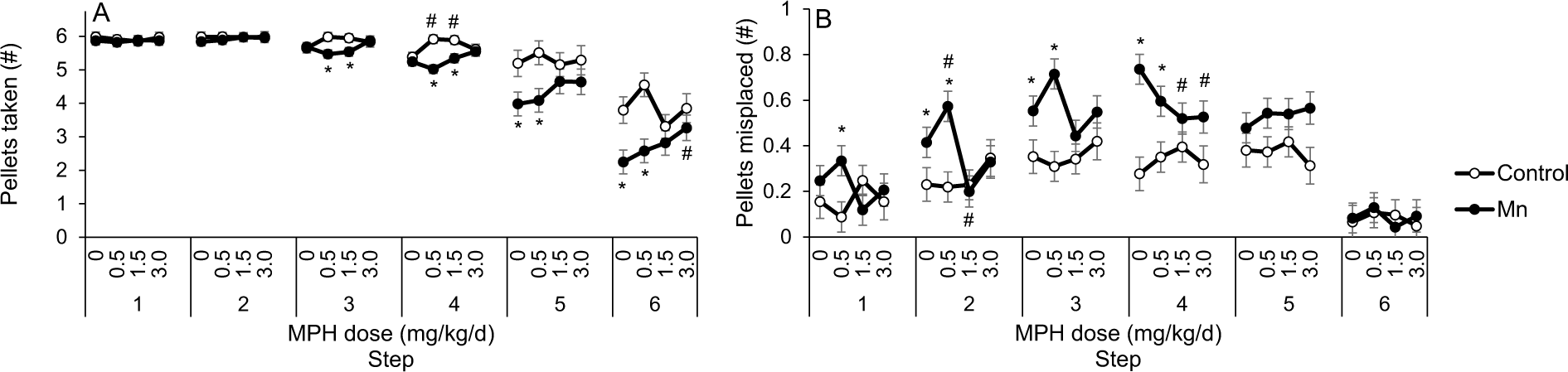
Developmental postnatal Mn exposure causes lasting sensorimotor deficits in the staircase test, and methylphenidate treatment at the highest 3.0 mg/kg/d dose alleviates those deficits. (A) Pellets taken (#) and (B) pellets misplaced (#) for the control and the Mn groups, as a function of MPH dose and staircase step. Data are lsmeans ± SEM of the control and Mn groups (*n* = 14–15/group). *p≤0.05 versus controls, and ^#^p≤0.05 versus 0 mg/kg/d MPH dose.

Similarly, the significant Mn x MPH x step interaction for the number of pellets misplaced reflects a Mn-induced impairment in sensorimotor grasping skills, and the efficacy of the higher MPH doses to alleviate this dysfunction. Specifically, this 3-way interaction reflects that the Mn+0 MPH group misplaced significantly more pellets than the Control+0 MPH group from staircase steps 2, 3 and 4 (p’s=0.030, 0.013, and 0.0001, respectively), and that this deficit was also ameliorated by the 1.5 and 3.0 MPH doses (Figure 5B).This is supported by contrasts showing that the Mn+1.5 MPH group misplaced significantly fewer pellets than the Mn+0 MPH group from steps 2 and 4 (p=0.008 and 0.050), with a similar trend seen in the 3.0 mg MPH/kg/d group from step 4 (p=0.069) (Figure 5B). In contrast, the 0.5 mg MPH/kg/d showed little or no efficacy to alleviate this sensorimotor deficit, in that the number of pellets misplaced by the Mn+0.5 MPH group was not significantly different than the Mn+0 MPH group from steps 2-4 (p’s=0.49, 0.14, and 0.20) (Figure 5B).

Finally, we performed follow-up day-by-day analyses to gain insight into whether prolonged treatment was needed to produce the benefits of MPH on sensorimotor function. For this, we focused analyses on the staircase steps where the Mn deficits and MPH efficacy were most apparent. These analyses revealed that the efficacy of the 1.5 or 3.0 MPH doses to ameliorate the Mn sensorimotor deficits was apparent early in treatment (e.g., after day 1), in contrast to the efficacy of the 0.5 MPH dose to fully ameliorate the Mn attention deficit, which only emerged after prolonged treatment (see Supplemental Material, *Figure S3A-D Montoya staircase follow-up analyses: Temporal trends in performance with methylphenidate treatment*).

### 3.5. Specific D1R and D2R involvement in Mn-induced sensorimotor impairments and MPH effectiveness

#### 3.5.1. The Mn-induced sensorimotor impairments and MPH efficacy persist throughout the antagonist treatment phase of the study

We first sought to confirm that the Mn-induced dysfunction in reaching and grasping, and the efficacy of the 3.0 mg MPH/kg/d treatment, persisted into the 36-day receptor antagonist phase of the study. For this, we analyzed the number of pellets taken and the number of pellets misplaced in the control/Mn groups treated with MPH under the antagonist treatment vehicle, which revealed a significant Mn x MPH x step interaction for the number of pellets taken and the number of pellets misplaced (F(15, 1419)=1.78, p=0.032, and F(15, 1453)=3.25, p<0.0001, respectively). The 3-way interaction for pellets taken reflects that the Mn+0 MPH group took significantly fewer pellets than their Control+0 MPH counterparts at staircase steps 4-6 (p’s=0.002, 0.006, and 0.005). Further, the interaction reflects the continued efficacy of MPH treatment; the Mn+1.5 MPH group took significantly more pellets than the Mn+0 MPH group at step 5 (p=0.046), and the Mn+ 3.0 MPH group took significant more pellets than the Mn+0 MPH group at step 6 (p=0.019, with a similar trend seen at step 5, p=0.094) (Figure 6A), consistent with our findings in the preceding phase of the study without receptor antagonists (described above). This confirms that the Mn *grasping deficit* and MPH efficacy persisted into the receptor antagonist phase of the study. This is further supported by the fact that the Mn+1.5 MPH group misplaced significantly fewer pellets than the Mn+0 MPH group from steps 1 and 4 (p’s=0.030 and 0.002), with a similar pattern of MPH efficacy seen at the 3.0 mg MPH/kg/d dose from steps 1, 2, and 4 (p=0.003, 0.019, and 0.046, respectively) (Figure 6B).

**Figure 6A-D.**
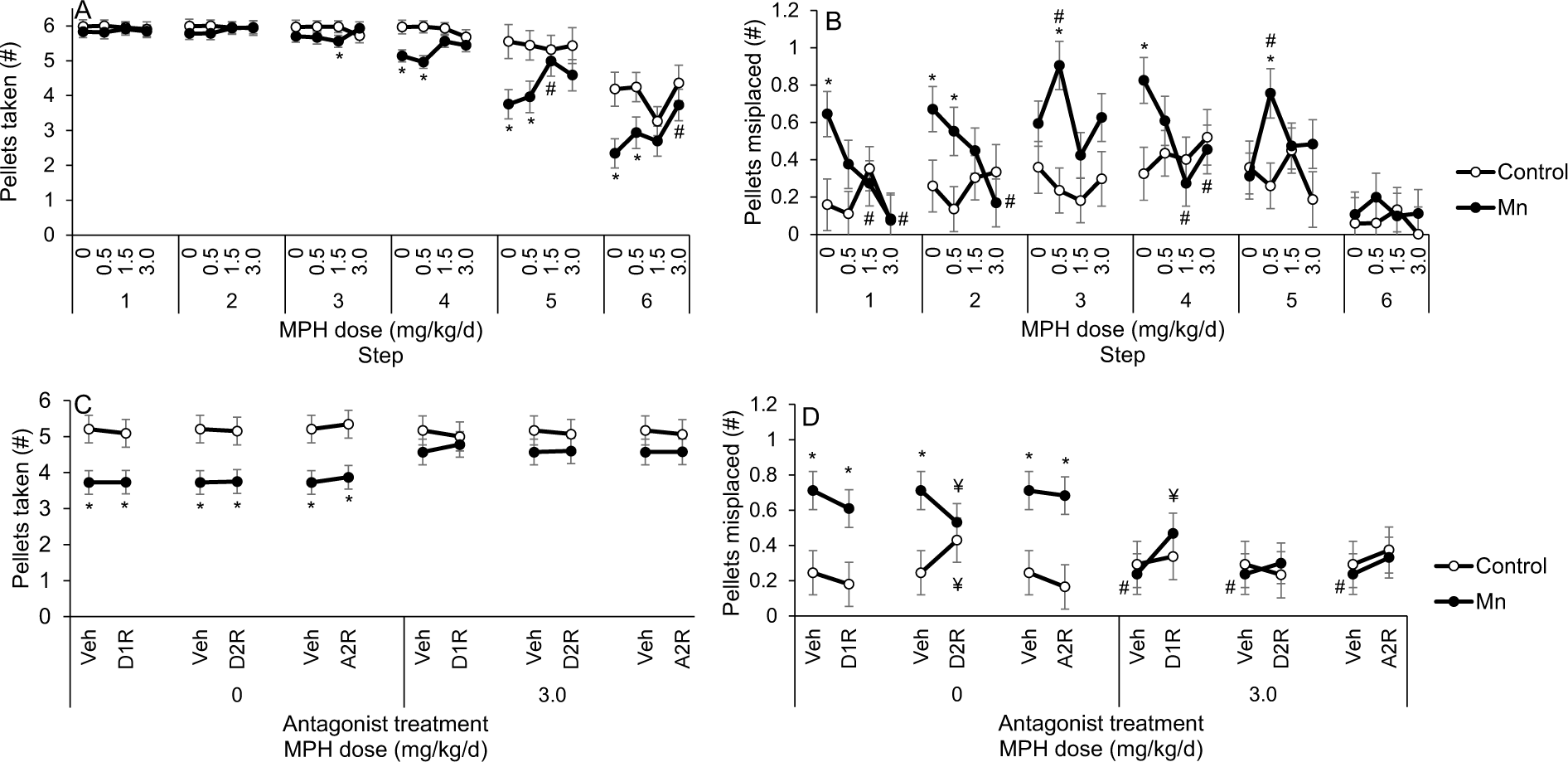
The Mn sensorimotor deficits and MPH efficacy to alleviate those deficits at the highest 3.0 mg/kg/d dose are sustained into the antagonist treatment phase in the staircase test, with D2R antagonism improving the Mn-induced grasping deficit in the absence of MPH, and D1R antagonism attenuating MPH efficacy to alleviate Mn grasping deficit at the 3.0 mg/kg/d dose. (A) Pellets taken (#) and (B) pellets misplaced (#) for the control and the Mn groups, as a function of MPH dose and staircase step. (C) Pellets taken (#) and pellets misplaced (D) for the control and the Mn groups, as a function of antagonist treatment and MPH dose. Data are lsmeans ± SEM of the control and Mn groups (*n* = 14–15/group). *p≤0.05 versus controls, ^#^p≤0.05 versus 0 mg/kg/d MPH dose, and ^¥^*p*≤0.05 versus vehicle antagonist treatment. Data in A and B are lsmeans ± SEM of the control and Mn-exposed groups under the vehicle antagonist treatment condition. The statistical model for C and D included, respectively, staircase step data 4-6 or 1-4 where there was an apparent effect of Mn exposure and/or MPH treatment.

#### 3.5.2. None of the specific receptor antagonists we studied directly mediate either the Mn reaching deficit or the efficacy of MPH to alleviate this deficit

Given the lasting reaching and grasping Mn deficits and efficacy of the 3.0 mg MPH/kg/d dose to alleviate these impairments at steps 4-6, we focused subsequent analyses regarding the effect of selective antagonist treatments on this MPH dose and these step levels. The analysis of the number of pellets taken revealed a trending 3-way interaction of Mn, MPH, and step (F(3, 116)=2.35, p=0.10), but no significant interactions between Mn and antagonist (F(3, 1478)=0.96, p=0.41), or between Mn, MPH, and antagonist (F(3, 1478)=0.25, p=0.86). These findings reflect that the Mn+0 MPH group took significantly fewer pellets than the Control+0 MPH under the vehicle, D1R, D2R, and α2_A_R antagonist treatment conditions (p=0.005, 0.009, 0.007, and 0.005, respectively) (Figure 6C). Additionally, the number of pellets taken by the Mn+3.0 MPH group was not different than that of the Control+0 MPH group across antagonist treatment conditions (p’s=0.23, 0.55, 0.30, and 0.15 for the vehicle, D1R, RD2R, and α2_A_R antagonist, respectively) (Figure 6C). We conclude from these results that the 3.0 mg/kg/d MPH dose ameliorated the sensorimotor Mn reaching deficit in pellets taken, and that none of the specific receptor antagonists singly mediate either the Mn reaching deficit or the efficacy of MPH to alleviate this deficit.

#### 3.5.3. The D2R antagonist ameliorates the sensorimotor Mn deficit in grasping skill, while the D1R antagonist partially blocks the efficacy of MPH therapy to ameliorate the Mn grasping deficit

The analysis of the number of pellets misplaced, a measure of sensorimotor grasping skill, revealed a significant Mn x MPH x antagonist interaction (F(3, 1478) = 3.81, p=0.009). This interaction reflects differential effects of Mn exposure, depending on the MPH and antagonist treatment conditions. Specifically, in the absence of MPH treatment, the selective D2R antagonist had opposing effects on the Mn and control groups, improving grasping abilities in the Mn group (i.e., indicated by the decreasing number of pellets misplaced; p=0.022), but impairing these abilities in the controls (indicated by a significant increase in the number of pellets misplaced (p=0.043), relative to antagonist vehicle treatment; Figure 6D). In contrast, treatment with the D1R and α2_A_R antagonists showed no measurable effect on the number of pellets misplaced (all p’s>0.40) in the Mn or control groups, compared to antagonist vehicle (Figure 6D).

Furthermore, the significant Mn x MPH x antagonist interaction for the number of pellets misplaced reported above indicates that the D1R antagonist attenuates the efficacy of MPH treatment to ameliorate the Mn-induced grasping deficit. Specifically, in the presence of MPH, treatment with the D1R antagonist substantially increased the number of pellets misplaced in the Mn group (p=0.007), but not in the controls (p=0.65). As a consequence of the D1R antagonist’s effect on the Mn+3.0 MPH group, there was no difference in the number of pellets misplaced between this group versus the Mn+0 MPH group treated with vehicle (p=0.13) or the D1R antagonist (p=0.37), demonstrating the re-emergence of the grasping Mn-induced impairment following D1R antagonism.

### 3.6. Postnatal Mn exposure produced environmentally relevant body Mn levels, with no effects on blood hematocrit

Oral exposure to 50 mg Mn/kg/d over PND 1 – 21 increased blood Mn levels in PND 24 littermates similar to those reported in our previous studies (Beaudin et al., 2017a; Beaudin et al., 2013), without impacting hematocrit levels in the same PND 24 littermates (see Supplemental Material, *Oral exposure producing environmentally relevant body Mn levels*).

## 4. DISCUSSION

Epidemiological studies have reported that environmental Mn exposure is associated with increased risk of ADHD and/or attention/impulsivity/hyperactivity symptoms in children and adolescents (Bhang et al., 2013; Karatela et al., 2022; Liu et al., 2020; Lucchini et al., 2012; Schullehner et al., 2020), although these studies were not able to determine causality, elucidate underlying mechanisms, or identify effective treatments. Here, using an animal model, we demonstrate that early developmental Mn exposure can *cause* lasting impairments in attention, learning, and sensorimotor functions, and that oral MPH treatment is effective in lessening all of these impairments, although the most efficacious MPH dose varies by area of dysfunction. Finally, we found that antagonism of specific D1, D2, or α2_A_ receptors does not alter either the degree of attentional impairment caused by developmental Mn exposure, or the MPH efficacy to improve Mn-induced attentional dysfunction, indicating that none of these receptor types singly mediates the Mn attentional dysfunction or MPH efficacy in this domain. In contrast, antagonism of D2R lessened Mn deficits in sensorimotor grasping skills in the absence of MPH, while antagonism of D1R attenuated MPH efficacy to ameliorate those deficits. These findings significantly advance understanding of the relationship between developmental Mn exposure and lasting attentional and sensorimotor deficits and their underlying mechanisms, and they demonstrate that a clinically-relevant oral MPH regimen is fully efficacious in ameliorating these deficits. The basis for each of these conclusions is provided below.

### 4.1. Mn-induced neurobehavioral deficits and efficacy of oral MPH treatment

A short period of early postnatal Mn exposure produced lasting impairments in learning, attention, and sensorimotor function, paralleling the domains of dysfunction seen in children exposed to elevated Mn levels (Bhang et al., 2013; Carvalho et al., 2018; Karatela et al., 2022; Lucchini et al., 2012; Oulhote et al., 2015; Schullehner et al., 2020). Notably, these impairments lasted well beyond the period of developmental Mn exposure, indicative of persistent or irreversible neurologic dysfunction due to the early exposure. For the four behavioral tasks administered here, the Mn group, in the absence of MPH, consistently responded less accurately to brief visual cues than controls, indicating attentional dysfunction due to early Mn exposure, as previously reported (Beaudin et al., 2017a). Importantly, treatment with the lowest dose of oral MPH (0.5 mg/kg/d) completely ameliorated the attentional dysfunction of the Mn animals, though this benefit only arose after prolonged MPH treatment (>∼9 days).

The fact that the efficacy of this low MPH dose was seen only after prolonged, but not acute, MPH treatment suggests that the efficacy is not strictly due to the acute pharmacologic action of the drug to increase synaptic dopamine and norepinephrine levels but may also be due to adaptive neuromolecular changes that emerge with prolonged treatment. Consistent with this inference, Yde Ohki et al. (2020) present evidence that chronic MPH treatment alters a number of intracellular signaling pathways, including Wnt- and mTOR signaling, with downstream changes in catecholaminergic function. These actions of MPH, perhaps in addition to its pharmacologic effects as a DAT/NET antagonist to increase synaptic dopamine and norepinephrine levels, may account for MPH efficacy in the treatment of ADHD-related symptoms (Coghill et al., 2014; Faraone et al., 2021). Irrespective of the precise mechanisms, our findings demonstrate the effectiveness of prolonged low-oral dose MPH treatment to ameliorate the lasting attention deficits caused by developmental postnatal Mn exposure, consistent with clinical studies showing effectiveness of MPH in the treatment of ADHD symptoms (Coghill et al., 2014; Faraone et al., 2013).

Methylphenidate was also efficacious in ameliorating the Mn sensorimotor impairments seen in the Montoya staircase test, and this benefit arose early in treatment (i.e., after day 1). In this test, the Mn group (0 MPH) took fewer food pellets than did controls from the most challenging staircase steps, indicating impaired execution of forelimb sensorimotor function as a result of the developmental Mn exposure, as previously reported (Beaudin et al., 2013). Methylphenidate treatment improved these sensorimotor abilities in the Mn-exposed rats, but only at the highest (3.0 mg/kg/d) dose. Based on these findings, we may infer that the 30 days of oral 3.0 mg MPH/kg/d treatment altered the number and/or binding sensitivity of the D1 and D2 receptors in the Mn group, which have been shown to be downregulated and upregulated, respectively, within the fronto-cortico-striatal system of adult rats by the same developmental Mn exposure regimen used here (Conley et al., 2020; Kern et al., 2010; Kern & Smith, 2011). Finally, this therapeutic efficacy of MPH on sensorimotor function recapitulates our previous findings (Beaudin et al., 2015), and is consistent with clinical studies showing that MPH treatment can attenuate the manual dexterity deficits in children with ADHD-DCD (Bart et al., 2013; Brossard-Racine et al., 2015; Flapper et al., 2006; Soleimani et al., 2017).

There are some prior rodent findings which also indicate that the dose of MPH needed to improve functions mediated by the prefrontal cortex (such as attention) is lower than that needed to affect striatal-based functions (likely contributing to these sensorimotor abilities). Specifically, rodent studies have shown that acute low and clinically relevant MPH doses (e.g., 0.25 - 1.0 mg/kg) improved prefrontal cortex-dependent functions, such as attentional processes and working memory, by selectively increasing synaptic levels of dopamine and norepinephrine within the prefrontal cortex (Berridge et al., 2006; Schmeichel and Berridge, 2013; Spencer and Berridge, 2019). At relatively higher MPH doses, dopamine and norepinephrine levels are increased across numerous brain areas, including the striatum (Berridge et al., 2006; Schmeichel and Berridge, 2013), which is consistent with the relatively higher MPH doses needed to improve sensorimotor versus attentional function observed in the present study. Overall, the pattern of findings in clinical and animal model studies, which are corroborated by the present study, suggests that the most efficacious dose of oral MPH differs depending on the behavioral domain of function and/or the brain region(s) subserving those different functions.

### 4.2. Insights into the mechanisms underlying Mn neurobehavioral toxicity and MPH efficacy

There is compelling clinical evidence that ADHD-related symptoms are mediated, at least in part, by catecholaminergic system dysfunction in children (Arnsten, 2011, 2009; Del Campo et al., 2011; Faraone et al., 2021). Consistent with this, we have shown that developmental postnatal Mn exposure in rats, which causes lasting attentional and sensorimotor symptoms akin to ADHD/DCD, also causes lasting hypofunctioning of the cate-cholaminergic system, including reduced synaptic release of norepinephrine and dopamine, and changes in protein levels of tyrosine hydroxylase, D1R, D2R, and dopamine and norepinephrine transporters in fronto-cortico-striatal brain regions, but no changes in α2_A_ noradrenergic receptor (α2_A_R) levels (Conley et al., 2020; Kern & Smith, 2011; Lasley et al., 2020). However, the receptor-specific antagonism findings from the present study suggest that neither D1R, D2R, or α2_A_R are singly responsible for the lasting Mn deficits in attention and sensorimotor reaching function or the efficacy of MPH to ameliorate those deficits. The fact that these receptor-specific antagonists did not measurably alter the Mn attentional and reaching deficits or MPH efficacy in these domains cannot be attributed to using antagonist dose levels insufficient to antagonize the receptor and elicit a behavioral response, because there is clear evidence in the present study that antagonism of D1R and D2R did alter specific behaviors, including inhibitory control of premature responses (Figure 5B), latency to initiate a correct response (Figure 5C), and grasping abilities (Figure 6). Collectively, these findings imply that numerous catecholaminergic and/or non-catecholaminergic receptor systems are involved in the Mn attention deficit and MPH effectiveness. This conclusion is consistent with research demonstrating that both ADHD symptoms and their effective treatment include several receptor systems (e.g., catecholaminergic, cholinergic, GABAergic, and glutamatergic) (Chan et al., 2016; Del Campo et al., 2011; Gamo and Arnsten, 2011; Huss et al., 2016; Purkayastha et al., 2015; Rizzo and Martino, 2015; Wigal et al., 2010).

In contrast to the null findings on the effects of the receptor antagonists on attentional function, the D1R-specific antagonist, but not the D2R or α2_A_R antagonists, reduced the effectiveness of the 3.0 mg/kg/d MPH dose in alleviating the sensorimotor grasping impairment caused by development Mn exposure. Specifically, the Mn+3.0 MPH+Veh animals performed significantly better than their Mn+0 MPH+Veh counterparts, and this improved sensorimotor performance in the Mn+3.0 MPH +Veh group was attenuated by the D1R-specific antagonist (SCH 23390) (i.e., in the Mn+3.0 MPH+D1R group, Figure 6D). These findings indicate that the D1R plays an important role in MPH efficacy to improve the lasting sensorimotor grasping deficits caused by developmental Mn exposure.

Interestingly, in the absence of MPH, antagonism of D2R with raclopride ameliorates the sensorimotor grasping dysfunction caused by Mn (Figure 6D), demonstrating that this domain of sensorimotor Mn deficits is mediated directly or indirectly by D2 receptor activity. This inference is supported by previous findings that the developmental Mn exposure regimen used here leads to lasting increases in D2R levels in the prefrontal cortex (Conley et al., 2020; Kern & Smith, 2011), as well as findings in the current study that antagonism of D2R impairs rather than improves grasping abilities in control rats (Figure 6D). Combined, these findings suggest that D2R may play a role in sensorimotor impairments caused by Mn possibly via the dopaminergic mesocortical system.

### 4.3. Strengths and limitations

There are several notable strengths of the present study. First, the nature of the behavioral tests allowed us to uncover specific cognitive and sensorimotor impairments produced by Mn – something that is often not possible to establish with the tasks used in human epidemiological studies. Specifically, the attention tasks included varying cue delays and durations as well as distraction conditions to create trials with varying demands on different aspects of attention. Similarly, the Montoya staircase task included steps which place increasing demands on sensorimotor grasping and reaching. Thus, in these tasks, the specific conditions under which the Mn animals differ from controls permits quite specific inferences about the nature of the impairments. Second, because our rodent model of Mn exposure recapitulated the areas of behavioral dysfunction reported in children and adolescents exposed to elevated levels of Mn, we had a sensitive animal model to test potential therapies and elucidate underlying neural mechanisms. Third, we utilized multiple prolonged oral doses of MPH, at clinically relevant doses. Finally, we employed specific catecholaminergic receptor antagonists to assess the neuromechanisms underlying the Mn effects and MPH efficacy.

There are also several limitations to our study. First, our conclusion that the optimal MPH dose for ameliorating the attentional dysfunction is lower than that needed for improving sensorimotor dysfunction must be tempered by the fact that the time interval between the daily oral MPH dosing and behavioral testing was different for the attention task and sensorimotor testing (∼30 and 90 min, respectively), possibly leading to different MPH pharmacokinetic profiles and brain MPH levels during these two tasks. Similarly, the receptor antagonism findings should be viewed within the context of their receptor binding properties (see Supplemental Materials: *Catecholaminergic receptor antagonist treatment*) for example, the apparent absence of D2R involvement in MPH efficacy may be a consequence of the D2R antagonist (raclopride) being displaced by dopamine under MPH treatment, which may have negated the antagonist effect on the MPH response of the Mn group.

## 5. CONCLUSIONS

Our findings demonstrate that developmental postnatal Mn exposure causes long-lasting attentional and sensorimotor dysfunction in adulthood, and that oral MPH treatment fully ameliorates those Mn deficits, although the most efficacious MPH dose varied for the different domains of neurobehavioral impairment. Notably, the MPH benefit on attention was only apparent after prolonged treatment, while MPH efficacy for the sensorimotor deficits emerged early in treatment. Further, selectively antagonizing D1, D2, or α2_A_ receptors had no effect on the Mn-induced attentional deficits, or MPH efficacy to ameliorate those deficits. In contrast, antagonism of D2R attenuated the sensorimotor deficits produced by Mn, whereas the efficacy of MPH to ameliorate those sensorimotor deficits was diminished by D1R antagonism. Collectively, these findings have significant implications for the mechanistic understanding and therapeutic treatment of environmentally-induced attentional and psychomotor dysfunction in children.

## ACKNOWLEDGEMENTS

We thank Shannon Twardy, Vanessa Howland, Gladys Chiang, Bhatia Bilow, Tyler Waite, Ramtin Poustinchi, Nelida Robles-Ramirez, Jonathan Do, Victoria Deng, Ayushi Modi, Naomi Nakamura, Dennis Luu, Ana Bayliss Grijalva, Roshana Basto, Levon Shahnazarin, Charlyn Agbayani, Aleksandra Nikolaeva, Ronald Lingat, Mariana Prado Martinez, Elizabeth Ngo, Aliyah Peterson, Yash Hothi, Jaelene Tapia, Wendolyne Valdez Rodriquez, and Jordan Altshul for assistance with the attention and sensorimotor series testing, and Tom Jursa for tissue analyses for Mn concentrations. Funding was from the National Institutes of Environmental Health Sciences (#R01ES028369).

## AUTHOR CONTRIBUTIONS

D.R.S., S.A.B. and B.J.S. conceived and designed research. S.A.B, S.H., N.S., and D.R.S. performed experiments. S.A.B., B.J.S. and D.R.S. analyzed and interpreted results of experiments. S.A.B. prepared figures. S.A.B., B.J.S. and D.R.S. drafted manuscript. S.A.B., B.J.S. and D.R.S. edited and revised manuscript.

