## Supplemental for "Methylphenidate alleviates cognitive dysfunction from early Mn exposure: Role of catecholaminergic receptors"

1. Subjects housing, feeding, and accreditation
2. Manganese exposure protocol and rationale
3. Methylphenidate treatment and rationale
4. Catecholaminergic receptor antagonist treatment
5. Behavioral testing procedures
6. 5-CSRTT dependent measures
7. Staircase test dependent measures
8. Montoya staircase follow-up analyses: Assessment of MPH efficacy as a function of days of treatment
9. Methods for determining blood Mn concentrations and hematocrit
10. Oral exposure producing environmentally relevant body Mn levels
11. References cited in Supplemental Material

Figure S1. Schematic diagram of study design and key elements of the behavioral study.

Figure S2A, B. Methylphenidate (MPH) dose-response for %accurate and %premature responses in the third focused attention task.

Figure S3A-D. Day-by-day sensorimotor performance with methylphenidate treatment in the Montoya staircase test.

**Supplemental material**

### 1. Subjects housing, feeding, and accreditation

A total of 128 male Long-Evans rats were used for neurobehavioral evaluation, and an additional 52 male and female littermates were used for blood Mn and hematocrit analyses. Animals were born from 28 nulliparous timed-pregnant rats (Charles River; gestational age 16). Twelve to 24 hours after parturition (designated PND 1, birth = PND 0), each litter was culled to eight pups, with five to six males and the remainder females. Only one male/litter was assigned to a particular treatment condition.

Animals (dams and weaned pups) were fed Harlan Teklad rodent chow #2018 (reported by the manufacturer to contain 118 mg Mn/kg) and housed in polycarbonate cages at a constant temperature of 21 ± 2°C. At weaning on PND 22, weanlings were pair-housed by treatment group assignment and kept on a reversed 10:14 hrs light/dark cycle. All aspects of testing, feeding, and treatment were conducted during the active (dark) phase of the cycle, approved by the institutional IACUC and adhered to NIH guidelines set forth in the Guide for the Care and Use of Laboratory Animals (NRC, 2011).

### 2. Manganese exposure protocol and rationale

Neonates were orally exposed to 0 or 50 mg Mn/kg/d over PND 1 - 21 (starting ~24 hrs after birth, designated PND 0). This Mn exposure level is not overtly toxic and does not measurably affect neonate health or nutrition, based on neonate milk intake from the lactating dam, growth rate, or blood hematocrit levels at weaning (Beaudin et al., 2017a; Beaudin et al., 2013; Kern et al., 2010). For Mn dosing, a 193 mg Mn/mL stock solution was prepared by dissolving MnCl2·4H2O with Milli-Q™ water; aliquots of the stock solution were diluted with a 2.5% (w/v) solution of the natural sweetener stevia for oral dosing of the neonates. Dosing solutions ranged from 0.02 to 0.097 mg Mn/µL over the course of pre-weaning exposure as the animals gained body wt. Doses were delivered once a day directly into the mouth of the pups in a volume of ~10 - 25 µL/dose via a micropipette fitted with a flexible polyethylene gel loading tip (Fisher Scientific, Santa Clara, CA, USA). Control animals received only the Milli-Q water + stevia vehicle.

The environmental relevance of the oral Mn dosing regimen used here is based on the following, as previously provided (Beaudin et al., 2017a; Kern et al., 2010): The 50 mg Mn/kg/d exposure levels over the pre-weaning period produce relative increases in Mn intake that are ~700-fold over levels consumed from lactation alone, which approximates the relative ~300 to 500-fold increases in Mn exposure experienced by infants and young children exposed to Mn-contaminated water or soy-based formulas (or both), compared to Mn ingestion from human breast milk (Kern et al., 2010). For example, human breast milk contains ~6 µg Mn/L, yielding normal infant intake rates of ~0.6 µg Mn/kg/d, based on infant daily milk consumption rates of ~0.8 L/day for a 8 kg 6-9 month old infant . Infants consuming contaminated water (e.g., directly or indirectly to rehydrate powdered formulas) containing 1.5 mg Mn/L, i.e., a level three-times the maximum contaminant level (MCL) guideline and comparable to median well-water levels associated with cognitive deficits and other effects in children, would experience Mn exposure of ~200 µg/kg/d, which is ~300- fold higher than the level of Mn intake from breast milk based on median fluid intake rates of ~1 L/d for infants. Our lab and others have shown that this, and similar, oral Mn exposure regimens impaired attention, learning, impulse control, locomotor activity, and sensorimotor function, and altered the fronto-striatal catecholamine system in adult rats (Beaudin et al., 2017a; Beaudin et al., 2013; Conley et al., 2020; Kern & Smith, 2011; Kern et al., 2010; Lasley et al., 2020; McDougall et al., 2008; Reichel et al., 2006).

#### 3. Methylphenidate treatment and rationale

To mimic clinical oral use of methylphenidate, the drug was delivered orally using cookie wafers (Mini-Vanilla, Nabisco Inc.), as described previously by us and others (Beaudin et al., 2017b; Beaudin et al., 2015; Ferguson & Boctor, 2009). Briefly, cookie wafers (Mini-Vanilla, Nabisco Inc) were quartered and treated with vehicle or MPH solution (~20–40 µL) to achieve the targeted daily drug dose per animal. The MPH solution was prepared fresh daily by dissolving the drug in normal saline solution. Adulterated pieces of wafer were transferred manually into individual food cups, placed onto the floor of the animal's individual holding cage, and typically consumed within 10 sec of delivery, which was confirmed by research personal delivering the pieces of wafer.

The clinical relevance of the oral MPH treatment regimen administered here is based on the following: the oral 0.5, 1.5, or 3.0 MPH mg/kg/d (or similar) MPH dosing regimens have been showed to be safe to adolescent rats and juvenile monkeys, to produce a blood MPH half-life of ~2 h in adult rats, and to improve attention, working memory, inhibitory control, and forelimb skill deficits in animal models with attention/hyperactivity deficits, as well as healthy animal subjects (Beaudin et al., 2017b; Beaudin et al., 2015; Cao et al., 2012; Kuczenski & Segal, 2002; Mohamed et al., 2011; Thanos et al., 2015; Zhu et al., 2007).

#### 4. Catecholamine receptor antagonist treatment

In the antagonist Phase 3 of the study, oral MPH treatment continued while animals also received a subcutaneous injection (1 mL/kg) of one of the catecholaminergic receptor antagonists (SCH23390 for D1R, raclopride for D2R, or BRL 44408 for $\alpha 2_A$R), or vehicle (saline) 10 minutes before MPH administration and 40 minutes before testing on the selective attention task, and ~100 min before the staircase task. Antagonist treatments were delivered at doses of 0.005 mg SCH23390 /kg s.c., 0.015 mg raclopride/kg s.c., and 1.0 mg BRL 44408/kg s.c. according to a within-subject quasi Latin-square design, in which all rats received all four antagonist treatments in each of three treatment cycles (i.e., three times). To minimize any potential antagonist treatment carry-over effect within a treatment cycle, the antagonist treatments were separated by 2-day washout breaks, with the first washout day consisting of no MPH and antagonist drug treatment and no testing, and the second washout day consisting of MPH treatment only (no antagonist) and testing on the third focused attention task. Each of the three treatment cycles (four antagonist treatment days plus 2 washout days per antagonist) lasted 12 days, and the entire antagonist treatment phase lasted 36 days (PND 144 to 179).

Over the receptor antagonist phase of the study, animals were treated with specific D1, D2 or $\alpha 2_A$ receptor antagonists (or vehicle) in the presence of continued MPH treatment, with testing first on the selective attention task followed by testing on the sensorimotor staircase task. Antagonists were administered 40 min before the start of behavioral testing, and 10 min prior to oral MPH dosing for the selective attention task. Receptor antagonist treatments were delivered according to a within-subject quasi Latin-square design, in which all subjects received the four different antagonist treatments over three consecutive treatment cycles, each lasting 12 days, for a total of 36 days (from ~PND 144 to 179). For each treatment cycle, the order of receptor antagonist treatment was completely balanced between the eight Mn and MPH treatment groups so that an equal number of animals in each treatment group received the four different receptor antagonist treatments in four different orders. For this, the order of receptor antagonist treatment was assigned pseudo-randomly, corresponding to four letters A-D, with each letter determining the first receptor antagonist treatment and dictating the order of the remaining three receptor antagonist treatments for a given animal and cycle. For instance, any animal in one of the eight treatment groups that was pseudo-randomly assigned the letter A first received the vehicle receptor antagonist treatment, followed by the D1, the D2, and the A2 receptor antagonist treatments. Another animal pseudo-randomly assigned the letter B received the four receptor antagonist treatment in a different order, starting with the D1 receptor antagonist treatment followed by the A2, the Veh, and the D2 receptor antagonist treatments. Pseudo-random assignment to the letters C and D delivered the four receptor antagonist treatment in yet different orders, corresponding to D2, Veh, A2, and D1, or A2, D2, D1, and Veh, respectively. This procedure allowed for the control of the order effect of each receptor antagonist treatment and yielded a total of 3 days of behavioral testing on the selective attention task and staircase test in the presence of MPH and each of the specific receptor antagonists. Furthermore, to minimize any potential antagonist treatment carry-over effect within a treatment cycle, the antagonist treatments were separated by 2-day washout breaks, with the first washout day consisting of no MPH and antagonist drug treatment and no testing, and the second washout day consisting of MPH treatment only (no antagonist) and testing on the third focused attention task. By the start of the antagonist treatment phase, the animals had been treated with MPH for 30 days.

Receptor-specific antagonists were administered 1 mL/kg s.c. at doses of 0.005 mg/kg for SCH23390 (D1 antagonist), 0.015 mg/kg for raclopride (D2 antagonist), and 1.0 mg/kg for BRL 44408 ($\alpha 2_A$ antagonist). All receptor antagonists were purchased from Tocris Bioscience (Minneapolis, MN). Antagonists were prepared in sterile saline; SCH23390 and BRL 44408 readily dissolved, while full dissolution of raclopride was encouraged with solution warming and gentle sonication. All antagonist solutions were made fresh weekly, aliquoted and maintained frozen (-20 °C) until the day of use. The antagonists, doses, and timing of treatment were chosen based on their receptor-specific relative Ki's, which exceed by >100-fold the binding affinities of the receptors' natural ligands (dopamine and norepinephrine), and their well-defined pharmacokinetics (Alexoff et al., 2003; Alstrup et al., 2013; Hall et al., 1989; Han & Gu, 2006; Janhunen et al., 2011). Selection was also informed by previous research demonstrating that these drugs antagonize the effect of acute MPH treatment on behavioral and physiological measures of fronto-striatal functions relevant to ADHD in animal models (Arnsten &

Dudley, 2005; Bandyopadhyay et al., 2005; de Oliveira et al., 2006; Dwyer et al., 2010; Fan & Hess, 2007; Gamo et al., 2010; Granon et al., 2000; Gronier, 2011; Hemsley et al., 1999; Hoffman & Donovan, 1994; Janhunen et al., 2011; Kiss et al., 1995; Newman et al., 2008; Uzsoki et al., 2011).

Finally, because SCH23390 and raclopride have been shown to decrease the number of response trials in the 5-CSRTTT (Koskinen & Sirviö, 2001; Shoaib & Bizarro, 2005), the final dose selection for these drugs was based on the behavioral results of a pilot experiment. In the pilot study, the D1R, D2R, and $\alpha 2_A$R receptor antagonist treatments were given to control and Mn-exposed rats 40 minutes before behavioral assessment on the selective attention baseline task. The pilot study found that doses of 0.005 mg SCH23390/kg and 0.015 mg raclopride/kg had little to no effect on response rates when compared to vehicle, and notably higher response rates compared to the initially selected higher doses of 0.25 mg/kg and 0.50 mg/kg of the same drugs, respectively; the initial dose of 1.0 mg BRL 44408/kg, like lower doses, had no effect on response rate in animals. Based on these findings, dosages of 0.005 mg SCH23390/kg, 0.015 mg raclopride/kg, and 1.0 mg BRL 44408/kg were utilized in the receptor antagonist treatment phase of the study.

### 5. Behavioral testing procedures

All animals were tested in the 5-Choice Serial Reaction Time Task (5-CSRTT) and the Montoya staircase test to evaluate learning, attention, impulse control, and sensorimotor functions. Sixteen identical automated 5-CSRTT testing chambers fitted with odor delivery systems (#MED-NP5L-OLF, Med Associates, Inc., St. Albans, VT) were used, as described previously (Beaudin et al., 2017a). Behavioral training began on PND 47, with 1 week of food magazine and nose-poke training, followed by testing on two visual discrimination learning tasks with a fixed cue duration and no pre-cue delay, followed by a series of visual focused and selective attention tasks, as described below and previously (Beaudin et al., 2017a). During testing, all animals were kept on a modest food restriction schedule with water available *ad lib* throughout behavioral testing, as described previously (Beaudin et al., 2017a). Animals were tested 6 days/week during the active dark period of the diurnal cycle. For testing on the attention/impulse control tasks, a daily test session consisted of 150 trials or 50 min, whichever came first, depending on the test administered. Each trial was initiated by a nose-poke in the food magazine port, followed by a 3 sec turnaround time to allow the animal to reorient from the food magazine wall to the response wall; trial onset began after the 3 sec turnaround time. All behavioral testing was conducted by personnel blind to the treatment condition of the subjects.

Food magazine training: Each rat was first trained to retrieve the 45 mg food pellet rewards from the food magazine in the Med-Associates 5-CSRTT chambers. The first trial of a training session always began with a free-reward delivery and the illumination of the food magazine light and house light. A nose-poke to collect the free-reward extinguished the food magazine light. The house light remained illuminated. After 10 s had elapsed, the magazine light was illuminated again and another reward was delivered for the rat to collect. Magazine training continued in this fashion until each rat made 50 rewarded trials in a daily training session. No LEDs were illuminated on the opposite curved response wall during food magazine training. The order and time of testing were balanced by treatment and stayed the same each day for every rat throughout the different phases of behavioral evaluation.

Nose-poke training: Following food magazine training, each rat was trained to nose-poke into the five 2.5 x 2.5 cm response ports of the curved response wall of the chamber. Again, the first trial of a nose-poke training session always began with a free-reward delivery and the illumination of the food magazine light and house light. A nose-poke into the food magazine to collect the free reward extinguished the food magazine light. The house light remained illuminated. In stage 1 of nose-poke training, a nose-poke into any one of the five response ports resulted in the immediate delivery of a food reward and the illumination of the food magazine light. Nose-poke training continued in this fashion over five consecutive stages that rewarded nose-pokes to ports 1, 2, 3, 4, and 5, respectively. This was done to ensure that all response ports were equally associated with a food reward before the start of testing proper. Each stage was completed after 70 rewarded trials. No response port LEDs were illuminated during nose-poke training.

Visual discrimination task: Following food magazine and nose-poke training, animals were trained in the 5-choice visual discrimination task: The first trials of each testing session also always began with a free-reward delivery and the illumination of the food magazine and house lights. After the rat interrupted the photo beam at the entrance of the food magazine (and extinguished the food magazine light), it was given 3 s to turn and reorient towards the opposite response wall, where one of the five response port LEDs was illuminated until the rat made a nose-poke into a port or after 10 s had elapsed (i.e., limited time hold period), whichever came first. A nose-poke into the illuminated response port was tallied as a correct response and resulted in reward delivery and the illumination of the food magazine light. A nose-poke into a non-illuminated port was tallied as an incorrect response and was unrewarded and followed by a 5 s 'time-out' with the house light extinguished. Failure to make a nose-poke within the 10 s response interval (i.e., response omission) was also unrewarded and followed by a 5 s 'time-out' with the house light extinguished. The time-out was reset each time a response port nose-poke was recorded during the time-out period as a correction procedure. After the 5 s time-out, both the food magazine and house lights were illuminated to signal the start of the next trial. Each rat was tested on the learning task to a criterion of ≥80% correct responses over two out of three consecutive testing sessions.

*Focused attention tasks:* The first focused attention task used pre-cue delays of 0, 1, 2, or 3 sec and a fixed visual cue duration of 0.7 sec and was administered for 12 testing sessions from PND 80-93. The second focused attention task included longer pre-cue delays of 0, 3, 4, or 5 sec and variable visual cue durations of 0.4 or 0.7 sec and was administered for 12 daily testing sessions from PND 95-108. The animals were then tested on a third focused task, which included pre-cue delays of 3 or 4 sec and a fixed cue duration 0.7 sec; animals were tested in the third focused attention task for 18 testing sessions/days over PND 109 – 129 while receiving for the first time daily oral MPH treatment.

*Selective attention task with olfactory distractors:* The subsequent selective attention task was designed to evaluate the rats' ability to maintain a behavioral or cognitive set in the face of distracting olfactory stimuli (Beaudin et al., 2017a). The selective attention task included the same pre-cue delays and fixed cue duration used in the third focused attention task. In addition, on one third of the trials in each daily testing session, an olfactory distractor was presented from a non-cue response port 1–3 sec before presentation of the visual cue. Animals were assessed on the selective attention task for 12 testing sessions while they continued to receive daily oral MPH treatment over PND 130-143.

Olfactory distractors in the selective attention task: The nine different liquid odorants used as olfactory distractors were prepared from pure liquid extracts of anise, maple, almond, peppermint, rum, orange, butter, cinnamon, and coconut (McCormick & Company, Inc., MD USA). Different concentrations of the pure liquid extracts ranging from 2.5% to 10% (v/v) were diluted with propylene glycol in a final volume of 200 mL liquid odorant. Twenty-five mL of the final solution was transferred via pipette into individual odor delivery jars mounted on the olfactory module on the outside wall of the sound attenuating cubicle housing the 5-CSRTT chamber. Odorized air was delivered for ~1 s into a response port within the chamber, with the condition that the visual cue response port and olfactory distractor response port never coincided within a trial. Delivery of odorized air used air pumps and computer-controlled solenoid valves for each chamber. Each response port contained an air inlet port and air evacuation port. During odor delivery, a vacuum pump fitted to the air evacuation port within the response port evacuated the odorized air so that it remained within the confines of the response port and did not enter the greater testing chamber space.

### 6. 5-CSRTT outcome measures

The following formulas were used to calculate different performance measures in the series of attention tasks. The percent correct responses was calculated as the number of correct responses/(correct + incorrect + premature + omissions) × 100; the percent incorrect responses was calculated as above but with incorrect responses in the numerator; the percent accurate responses was calculated as the number of correct responses/(correct + incorrect) × 100; the percent premature responses was calculated as the number of premature responses/(correct + incorrect + premature + omissions) × 100; and the percent omission errors was calculated as the number of omissions/(correct + incorrect + premature + omissions) × 100. In addition, time latencies of nose-poke responses and latencies to collect food rewards following correct responses were also recorded on each trial in a session. Response latency was defined as the elapsed time between visual cue presentation and when the rat interrupted the photo beam at the entrance of one of the response ports. Food magazine latency was also recorded after each correct response and defined as the elapsed time between a response port nose-poke and when the rat interrupted the photo beam at the entrance of the food magazine.

### 7. Staircase test dependent measures

Step-by-step skilled forelimb performance was evaluated as previously described by us (Beaudin et al., 2015) and according to: the number of "pellets taken" per step describes those pellets that were grasped and removed from their step of origin and was calculated as [number of initial pellets - number of pellets remaining per step]. The number of "pellets eaten" describes pellets that were removed and consumed and was calculated as [number of initial pellets - (number of pellets remaining + number of pellets misplaced + the number of pellets lost)]; for this, the number of "pellets misplaced" describes those pellets that were grasped but ended up on a different step level than their initial starting placement, while the number of "pellets lost" describes those pellets that ended up on the floor of the apparatus out of reach of the animal. Finally, the "percent grasping/retrieval success" was calculated as the ratio of the number of pellets eaten/pellets taken x 100.

### 8. Montoya staircase follow-up analyses: Assessment of MPH efficacy as a function of days of treatment

We performed follow-up day-by-day analyses to assess whether MPH efficacy to ameliorate the Mn-induced sensorimotor dysfunction emerged early in treatment, or only emerged after prolonged treatment as with MPH efficacy to ameliorate the attentional deficits. For this, we focused analyses on the number of pellets taken and the number of pellets misplaced for each of the *first* 9 days of MPH treatment or the *last* 9 days of MPH treatment, focusing on staircase steps where the Mn-induced deficit was evident (i.e., steps 5 and 6 for pellets taken, and steps 3 and 4 for pellets misplaced).

Analyses uncovered a significant interaction of Mn, MPH, and day for the number of pellets misplaced for the analyses of both the *first* and the *last* 9 days of MPH treatment ($F(24, 1812)=1.57$, $p=0.039$ and $F(24, 1785)=1.61$, $p=0.031$, respectively), and the same 3-way interaction for the number of pellets taken over the *last* 9 days of MPH treatment ($F(24, 1784)=1.51$, $p=0.050$). There was no Mn x MPH x day interaction for the number of pellets taken over the first 9 days of MPH treatment ($F(24, 1751)=0.70$, $p=0.85$) (Figure S3), but there was a significant Mn x step interaction ($F(1, 1751)=6.56$, $p=0.011$), and a trending Mn x MPH interaction ($F(3, 104)=0.55$, $p=0.064$) for the number of pellets taken over the first 9 days of MPH treatment..

Specifically, for the number of pellets misplaced over the first 9 days of MPH treatment, the significant Mn x MPH x day interaction (reported above) reflects that grasping performance in the Mn+0 MPH group was significantly worse than the Control+0 MPH group on days 2, 3, and 6 (p's<0.033), and significantly worse in the Mn+0.5 MPH group than the Control+0.5 MPH group on days 3 and 4, 8 and 9 (p's≤0.05), while there were no significant differences in the number of pellets misplaced between the Mn+1.5 MPH and Control+1.5 MPH groups, or between the Mn+3.0 MPH and Control+3.0 MPH groups across days 1-9 of MPH treatment (p's>0.11). Collectively, these findings indicate that 1) on early testing days that showed a Mn grasping deficit there was complete efficacy of the 1.5 and 3.0 MPH doses, but not the 0.5 MPH dose, to ameliorate the Mn sensorimotor grasping deficit from the start of treatment, and 2) that MPH efficacy emerged very early in treatment (e.g., after day 1).

A qualitatively similar pattern of MPH efficacy was seen for sensorimotor reaching performance, as reflected in the significant Mn x step and trending Mn x MPH interactions for the number of pellets taken over the first 9 days of MPH treatment (Figure SX). Moreover, the significant Mn x MPH x day interaction over the last 9 days of MPH treatment reflects that reaching performance in the Mn+0 MPH group was significantly worse than the Control+0 MPH group on days 11 and 12 and 16 and 17 (p's<0.05), and significantly worse during almost all days in the Mn+0.5 MPH group versus their Control+0.5 MPH counterparts (all p's<0.03) (Figure S3). On the other hand, there were no significant differences in the number of pellets taken between the Mn+1.5 MPH and the Control+1.5 MPH groups, or between the Mn+3.0 MPH and the Control+3.0 MPH groups across days 10-18 of MPH treatment (p's>0.17). It is also noteworthy that, within each treatment group, the day-by-day reaching performance was relatively consistent over both the first and last 9 days of testing, except between days 15 and 16 under the vehicle condition (p=0.021); the latter finding explains the presence of the Mn x MPH x day interaction for the last 9 days of testing and the absence of an interaction involving day over the first 9 testing days. Collectively, these findings indicate that the effectiveness of MPH in ameliorating Mn-induced sensorimotor impairments emerges early on in treatment, unlike the efficacy of MPH to ameliorate the attention deficits caused by Mn exposure, which emerged only after prolonged MPH treatment.

### 9. Methods for determining blood Mn concentrations and hematocrit

Blood Mn concentrations were determined in littermates of the study animals at PND 24 and PND 66 (7 – 8/treatment group and time point). Animals were euthanized via sodium pentobarbital overdose (75 mg/kg i.p.) and exsanguination, and whole blood (2 – 3 mL) was collected from the left ventricle of the surgically-exposed heart and stored in EDTA Vacutainers at -20 ºC for analyses. Briefly, aliquots of whole blood were digested overnight at room temperature with 16N $HNO_3$ (Optima grade, Fisher Scientific), followed by addition of $H_2O_2$ and Milli-Q™ water. Digestates were centrifuged (15,000 x g for 15 min.) and the supernatant collected for Mn analysis. Rhodium was added to sample aliquots as an internal standard. Manganese levels were determined using a Thermo Element XR inductively coupled plasma – mass spectrometer, measuring masses $^{55}$Mn and $^{103}$Rh (the latter for internal standardization).  External standardization for Mn used certified SPEX standards (Spex Industries, Inc., Edison, NJ). National Institutes of Standards and Technology SRM 1577b (bovine liver) was used to evaluate procedural accuracy. The analytical detection limit for Mn in blood was 0.04 ng/mL.

For blood hematocrit, whole blood was collected into EDTA Vacutainers and hematocrit measured immediately using standard heparinized hematocrit capillary tubes and centrifugation (Kern et al., 2010).

### 10. Oral exposure producing environmentally relevant body Mn levels

Oral exposure to 50 mg Mn/kg/d over PND 1 – 21 increased blood Mn levels in PND 24, but not PND 66 animals. Specifically, blood Mn levels in Mn-exposed PND 24 littermates (385 ng/mL ± 55 SE, n=8) were significantly higher compared to control littermates (18 ng/mL ± 1.2) ($p<0.0001$), while in PND 66 littermates levels were notably lower and no longer different between control and Mn groups (13.5 ng/mL ± 1.2 SE vs 16.2 ± 1.3 for control and Mn groups, respectively) ($p>0.12$, n=7-8 per group). Also, blood hematocrit levels in PND 24 littermates were not different between control (40% ± 0.86 SE, n=16) and Mn (39% ± 0.75, n=21) groups ($p=0.343$).

| Phase | Mn exposure | Mn neurobehavioral evaluation | Mn + MPH treatment | Mn + MPH + Antagonist treatment |
|---|---|---|---|---|
| Factorial design | | One-way | 2 x 4 | 2 x 4 x 4 |
| Age (in PND) | 1 - 21 | 80 -108 | 109 -143 | 144 -179 |
| Duration (in total # of study day) | 21 | 24 | 30 | 36 |
| Treatment regimen | 0 and 50 mg Mn/kg/d | No treatment | 0, 0.5, 1.5 and 3.0 mg MPH/kg/d | 0, 0.5, 1.5 and 3.0 mg MPH/kg/d<br>Vehicle<br>0.005 mg SCH 23390/kg/d<br>0.015 mg raclopride/kg/d<br>1.0 mg BRL 44408/kg/d |
| Task administered | No testing | Focused attention 1 and 2 tasks with variable pre-cue delays | Focused attention 3 task, selective attention task with distractors, and staircase test | Selective attention task with distractors and staircase test |
| Function assessed | | Ability to maintain attention and inhibit response | Ability to maintain attention and behavioral set, inhibit response, and forelimb skills | Ability to maintain behavioral set, inhibit response, and forelimb skills |

Figure S1. Schematic diagram of study design and key elements of the behavioral study.

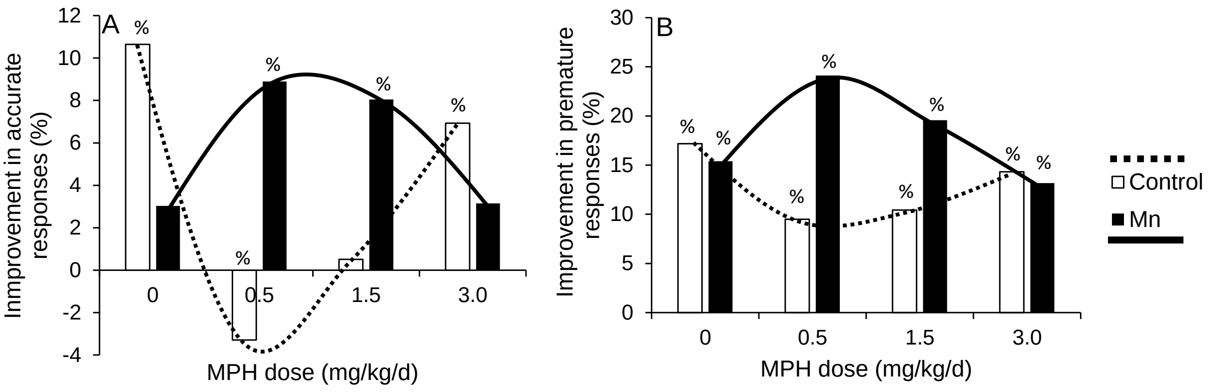


Figure S2. Methylphenidate (MPH) dose-response for %accurate and %premature responses in the third focused attention task. The MPH dose-response for attentional accuracy (A) and premature responses (B) are fundamentally different in the Mn (inverted U-shaped) compared to the control (U-shaped) groups. Improvement in percent accurate and premature responses in A and B was calculated for each Mn-exposed and MPH-treated groups by subtracting the group mean responses in session block 6 from the group mean responses in session block 1 (i.e., the delta). Improvement scores for premature responses are expressed positively. Higher scores reflect greater improvement in performance. % in panels A and B denotes percent accurate and premature responses, respectively, in manuscript Figures 2A and C session block 6 was different than block 1 at p≤0.05.

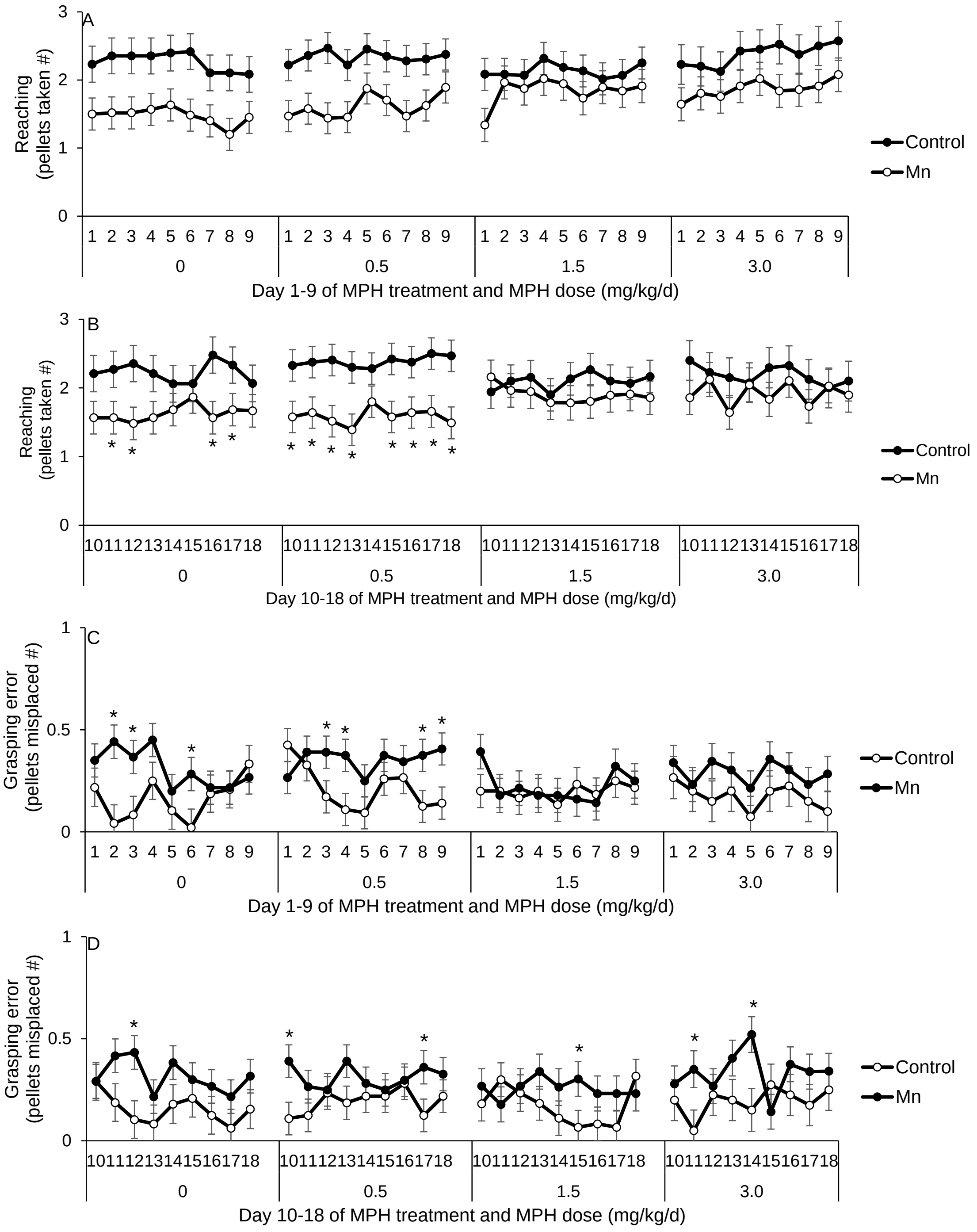


Figure S3. Day-by-day sensorimotor performance with methylphenidate treatment in the Montoya staircase test. Efficacy of the higher MPH doses to ameliorate Mn sensorimotor reaching and grasping dysfunctions emerged early in treatment. (A) Pellets taken over the first 9 days, and (B) the last 9 days of MPH treatment for the control and the Mn groups, as a function of MPH dose. (C) Pellets misplaced over the first 9 days, and (D) the last 9 days of MPH treatment for the control and the Mn groups, as a function of MPH dose. Data are lsmeans ± SEM of the control and Mn groups (*n* = 14–15/group). *$p \leq 0.05$ versus controls with each MPH dose treatment. The statistical model for A and B included staircase steps 5-6,and for C and D the staircase steps 3-4 where there was an apparent effect of Mn exposure and/or MPH treatment.
